# Demonstration of a common DPhe^7^ to DNal(2’)^7^ peptide ligand antagonist switch for the melanocortin-3 and melanocortin-4 receptors identifies systematic mischaracterization of the pharmacological properties of melanocortin peptides

**DOI:** 10.1101/2022.01.03.474807

**Authors:** Luis E Gimenez, Terry A. Noblin, Savannah Y. Williams, Satarupa Mullick Bagchi, Ren-Lei Ji, Ya-Xiong Tao, Claus B. Jeppesen, Kilian W. Conde-Frieboes, Tomi K. Sawyer, Paolo Grieco, Roger D. Cone

## Abstract

Melanocortin peptides containing a D-naphthylalanine residue in position 7 (DNal(2’)^7^), reported as melanocortin-3 receptor (MC3R) subtype-specific agonists in two separate publications, were found to lack significant MC3R agonist activity. The cell lines used at the University of Arizona for pharmacological characterization of these peptides, consisting of HEK293 cells stably transfected with human melanocortin receptor subtypes MC1R, MC3R, MC4R, or MC5R, were then obtained and characterized by quantitative PCR. While the MC1R cell line correctly expressed only the hMCR1, the three other cell lines were mischaracterized with regard to receptor subtype expression. Demonstration that a D-naphthylalanine residue in position 7, irrespective of the melanocortin peptide template, results primarily in antagonism of the MC3R and MC4R, then allowed us to search the published literature for additional errors. The erroneously characterized DNal(2’)^7^-containing peptides date back to 2003; thus, our analysis suggests that systematic mischaracterization of the pharmacological properties of melanocortin peptides occurred.

## Introduction

The discovery of linear and cyclic super-potent agonist analogs of the native melanocortin ligand α-melanocyte-stimulating hormone (α-MSH) ^1-3^ has led to the development of FDA-approved therapeutics for disorders as diverse as erythropoietic porphyria ^4^, syndromic obesity ^5^, and low libido ^6^. These basic principles, elucidated primarily by Dr. Victor Hruby and his colleagues, continue to guide the field today. However, the field has been challenged by the lack of receptor subtype-specific compounds.

Many G-protein-coupled receptors (GPCRs), such as the five melanocortin receptors, are members of receptor families, each activated by the same ligand or family of related ligands. Because each receptor subtype may play a unique physiological role, a critical goal of chemists and pharmacologists has been to design receptor-subtype specific ligands, often improving upon nature. In the case of the melanocortin receptors, this is important in that the five receptors each exhibit distinct sites of expression and regulate several unrelated physiological functions. The melanocortin-1 receptor (MC1R) is expressed in melanocytes and regulates eumelanin production in hair and skin ^7^, the MC2R is expressed in the adrenal cortex and regulates adrenal steroidogenesis ^7^, the MC3R and MC4R are primarily in CNS ^8-12^, where they regulate aspects of energy homeostasis ^13-15^, and the MC5R is expressed in exocrine glands, where it regulates the synthesis and secretion of exocrine gland products ^16^. The melanocortin therapeutics currently on the market lack receptor subtype specificity, a well-documented problem for these peptide drugs. For example, the drug Imcivree, an MC4R agonist used clinically to treat certain forms of syndromic obesity ^5, 17^, causes hyperpigmentation due to cross-reactivity with the MC1R in melanocytes ^18^.

Typically, cell lines transfected with expression vectors containing the cloned receptor subtype are used to characterize the receptor-subtype-specific pharmacology of existing or novel ligands. These lines also often include reporter systems to allow for facile production of concentration-response curves following treatment of cells with varying ligand concentrations under study. All five melanocortin receptors couple well to G_αs_ and the elevation of intracellular cAMP. Thus, in the case of the melanocortin receptors, these reporter systems have evolved, resulting in the production of multiple sets of different reporter cell lines, frequently in the HEK293 cell line. The initial characterization of the cloned receptors utilized a laborious biochemical method to quantify intracellular cAMP ^19^, followed by more facile cAMP RIA methods ^20^. These were further improved using a variety of academic or commercial systems, based on either gene expression ^21^ or enzymatic reporters of intracellular cAMP levels ^22^. These different reporter systems may yield different EC_50_ values for individual ligands, while properties such as the rank order of potency and agonist vs. antagonist activity remain unchanged.

Because melanocortin receptors all couple to G_αs_, and receptor-subtype reporter systems involve sets of five cell lines, often all in the HEK293 cell background, maintaining the identity and purity of these clonally-derived lines provides additional challenges. The misidentification of cell lines is a problem that has long been an issue in biomedical research ^23^. Based on research from institutional cell banks, up to 18% of lines submitted are misidentified ^24^. Mislabeling and cross-contamination are two leading causes of cell line misidentification. For example, a simple re-use of a pipette, along with different growth rates of clonal cell lines, can result in a contaminating cell overtaking a line in 4-5 passages. One of our laboratories (RDC) recently acquired melanocortin peptides PG-990 and PG-992, published as MC3R agonists from another author (PG) ^25^. Upon attempts at validating reported pharmacological properties in our laboratory (RDC), we could not repeat these findings. Further investigation determined that in vitro pharmacological characterization of these peptides had been performed by Dr. Minying Cai (MC) at University of Arizona. The data shown here demonstrate cell line misidentification at the University of Arizona to be a potential cause of the issue, identify published work that needs to be corrected, and provide a simple qPCR protocol for definitive characterization of human melanocortin receptor subtype-expressing cell lines to insure proper characterization of melanocortin peptides.

## Results

### Characterization of peptides PG-990 and PG-992

Published peptides reported to be MC3R specific agonists ^25^ were obtained (from PG, University of Naples, Table 1) by one of us (RDC, University of Michigan) for in vivo analysis of the physiological functions of the MC3R. These peptides were reported to be full agonists with 1.9nM (PG-990) and 42nM (PG-992) EC_50_ values at the human MC3R (hMC3R) while exhibiting no detectable agonist activity at the hMC4R at concentrations up to 1μM ^25^. Routine confirmational analysis of the activity of the peptides at the hMC3R and hMC4R was performed using clonal HEK293 cell lines constructed at the University of Michigan containing a cAMP split-luciferase reporter (Promega, Madison, WI), and individually expressing either hMC3R or hMC4R. Almost no agonist activity was detected at the hMC3R or hMC4R for PG-990 and PG-992 at peptide concentrations up to 10^−5^ M. (Figure 1A, C, Table 2). As controls, α-MSH and DTrp^8^-*γ*-MSH were also tested in parallel and exhibited EC_50_ values expected for these peptides, with DTrp^8^-*γ*-MSH exhibiting a ∼20-100X greater potency at hMC3R vs. hMC4R, as previously reported ^26^. Competition assays against an EC_80_-EC_90_ concentration of α-MSH were then performed, demonstrating that PG-990 and PG-992 are weak antagonists of the hMC3R and hMC4R (Figure 1B, D, and Table 3).

**Table 1.**
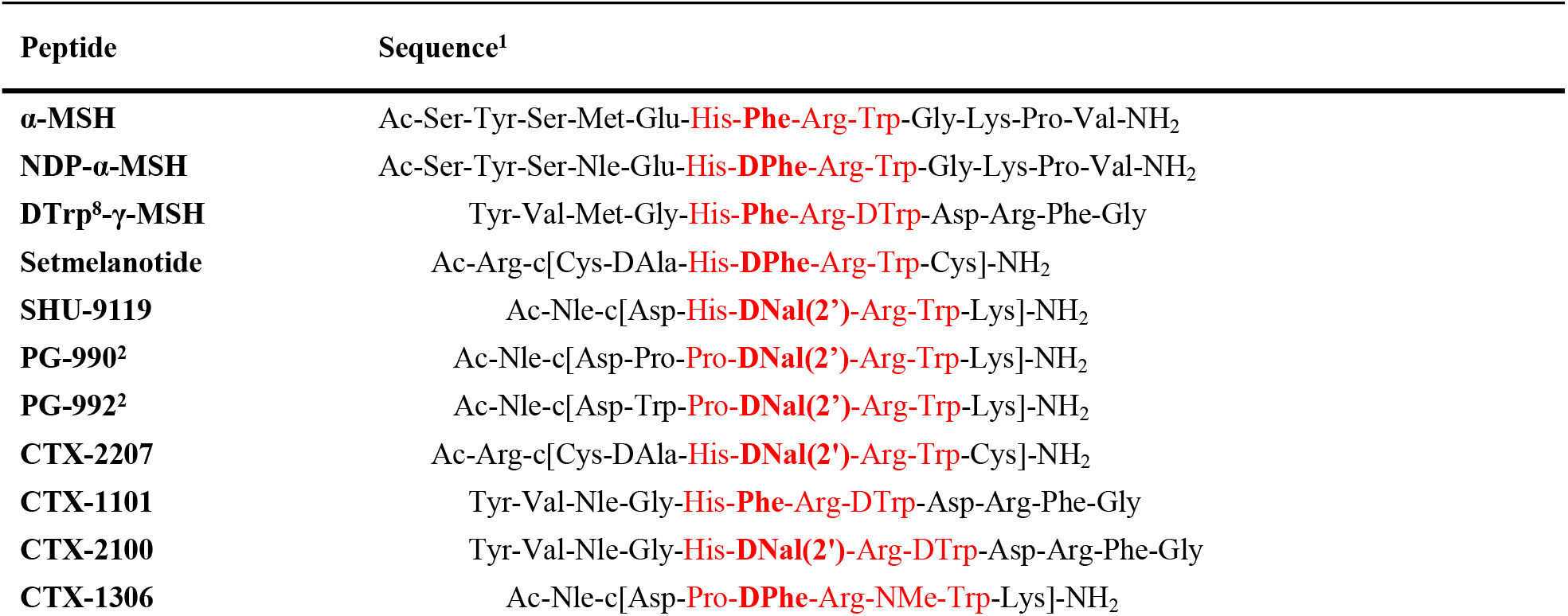
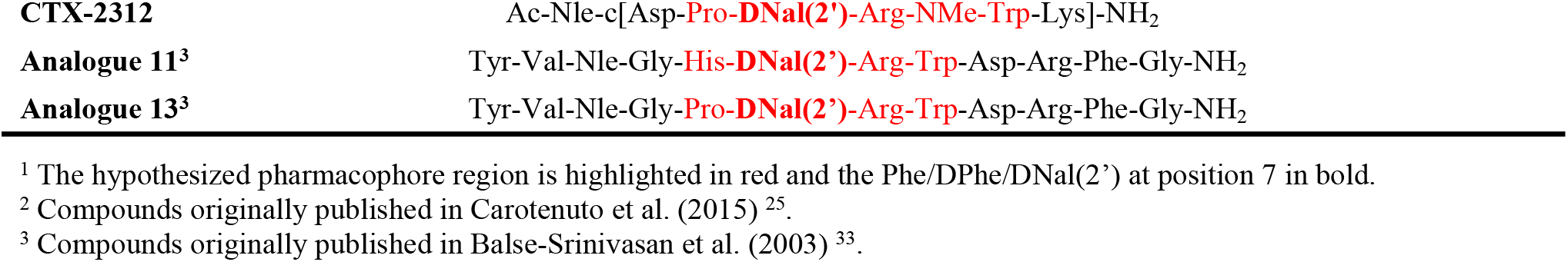
Sequences of peptides analyzed for this study.

| Peptide | Sequence<sup>1</sup> |
| --- | --- |
| α-MSH | Ac-Ser-Tyr-Ser-Met-Glu-His-Phe-Arg-Trp-Gly-Lys-Pro-Val-NH<sub>2</sub> |
| NDP-α-MSH | Ac-Ser-Tyr-Ser-Nle-Glu-His-DPhe-Arg-Trp-Gly-Lys-Pro-Val-NH<sub>2</sub> |
| DTrp<sup>8</sup>-γ-MSH | Tyr-Val-Met-Gly-His-Phe-Arg-DTrp-Asp-Arg-Phe-Gly |
| Setmelanotide | Ac-Arg-c[Cys-DAla-His-DPhe-Arg-Trp-Cys]-NH<sub>2</sub> |
| SHU-9119 | Ac-Nle-c[Asp-His-DNal(2')-Arg-Trp-Lys]-NH<sub>2</sub> |
| PG-990<sup>2</sup> | Ac-Nle-c[Asp-Pro-Pro-DNal(2')-Arg-Trp-Lys]-NH<sub>2</sub> |
| PG-992<sup>2</sup> | Ac-Nle-c[Asp-Trp-Pro-DNal(2')-Arg-Trp-Lys]-NH<sub>2</sub> |
| CTX-2207 | Ac-Arg-c[Cys-DAla-His-DNal(2')-Arg-Trp-Cys]-NH<sub>2</sub> |
| CTX-1101 | Tyr-Val-Nle-Gly-His-Phe-Arg-DTrp-Asp-Arg-Phe-Gly |
| CTX-2100 | Tyr-Val-Nle-Gly-His-DNal(2')-Arg-DTrp-Asp-Arg-Phe-Gly |
| CTX-1306 | Ac-Nle-c[Asp-Pro-DPhe-Arg-NMe-Trp-Lys]-NH<sub>2</sub> |
| CTX-2312 | Ac-Nle-c[Asp-**Pro-DNal(2')**-Arg-NMe-Trp-Lys]-NH<sub>2</sub> |
| Analogue 11<sup>3</sup> | Tyr-Val-Nle-Gly-**His-DNal(2')**-Arg-Trp-Asp-Arg-Phe-Gly-NH<sub>2</sub> |
| Analogue 13<sup>3</sup> | Tyr-Val-Nle-Gly-**Pro-DNal(2')**-Arg-Trp-Asp-Arg-Phe-Gly-NH<sub>2</sub> |

<sup>1</sup> The hypothesized pharmacophore region is highlighted in red and the Phe/DPhe/DNal(2') at position 7 in bold. <sup>2</sup> Compounds originally published in Carotenuto et al. (2015) <sup>25</sup>. <sup>3</sup> Compounds originally published in Balse-Srinivasan et al. (2003) <sup>33</sup>.

**Table 2.** Agonist pharmacological properties of PG-990, PG-992, analogue 11, and analogue 13 compared with published data.

| Data source | Compound | Receptor type |  |  |  |
| --- | --- | --- | --- | --- | --- |
|  |  | MC1R | MC3R | MC4R | MC5R |
| **Experimental potencies<sup>1</sup>** | **PG-990** | NA<sup>3</sup> | NA | NA | NA |
|  | **PG-992** | ND<sup>4</sup> | NA | NA | ND |
|  | **Analogue 11** | 9.79 ± 0.04 (**0.16**) | 8.21 ± 0.10 (**6.2**) | NA | 8.42 ± 0.21 (**3.8**) |
|  | **Analogue 13** | 8.91 ± 0.07 (**1.2**) | NA | NA | 8.38 ± 0.17 (**4.1**) |
| **Published potencies<sup>2</sup>** | **PG-990<sup>5</sup>** | 940 ± 100 | 1.9 ± 0.1 | > 1000 | 10.1 ± 0.1 |
|  | **PG-992<sup>5</sup>** | > 1000 | 42 ± 12 | > 1000 | 20 ± 4 |
|  | **Analogue 11<sup>6</sup>** | ND | > 10000 | 24 ± 2 | > 10000 |
|  | **Analogue 13<sup>6</sup>** | ND | 1700 ± 223 | 50 ± 5 | > 10000 |

<sup>1</sup> Experimental values obtained from split luciferase cAMP sensor dynamic assays are expressed as pEC<sub>50</sub> ± SEM and EC<sub>50</sub> values in parentheses (nM), and represent the mean from three independent experiments with five replicates each.
<sup>2</sup> Published values, from Carotenuto et al. (2015)<sup>25</sup> and Balse-Srinivasan et al. (2003)<sup>34</sup>, are expressed as EC<sub>50</sub> ± SEM in nM.
<sup>3</sup> NA; no activity at the compound concentrations assayed with a maximum efficacy ≤ 10% relative to α-MSH.
<sup>4</sup> ND; not determined.
Data on PG-990, PG-992, analogue 11, and analogue 13 were adapted from Carotenuto, A.; Merlino, F.; Cai, M.; Brancaccio, D.; Yousif, A. M.; Novellino, E.; Hruby, V. J.; Grieco, P. Discovery of novel potent and selective agonists at the melanocortin-3 receptor. *Journal of Medicinal Chemistry* 2015, 58, 9773-9778, and from Balse-Srinivasan, P.; Grieco, P.; Cai, M.; Trivedi, D.; Hruby, V. J. Structure-activity relationships of gamma-MSH analogues at the human melanocortin MC3, MC4, and MC5 receptors. Discovery of highly selective hMC3R, hMC4R, and hMC5R analogues. *Journal of Medicinal Chemistry* 2003, 46, 4965-4973, with permission from the *Journal of Medicinal Chemistry*, and the American Chemical Society.

**Table 3.** Antagonist pharmacological properties of PG-990, PG-992, analogue 11, and analogue 13.

| Data source | Compound | Receptor type |  |  |  |
| --- | --- | --- | --- | --- | --- |
|  |  | MC1R | MC3R | MC4R | MC5R |
| **Experimental potencies<sup>1</sup>** | **PG-990** | ND<sup>2</sup> | NC<sup>3</sup> | 5.96 ± 0.01 (**1096**) | ND |
|  | **PG-992** | ND | 5.67 ± 0.01 (**2149**) | 6.45 ± 0.02 (**568**) | ND |
|  | **Analogue 11** | 10.13 ± 0.05 (**0.74**) | 7.98 ± 0.13 (**10.6**) | 8.41 ± 0.23 (**3.9**) | NA<sup>4</sup> |
|  | **Analogue 13** | 8.20 ± 0.05 (**6.3**) | 7.99 ± 0.07 (**10.2**) | 8.40 ± 0.19 (**4.0**) | NA |

<sup>1</sup> Experimental values obtained from split luciferase cAMP sensor dynamic assays are expressed as pIC<sub>50</sub> ± SEM and IC<sub>50</sub> values in parenthesis (nM), and represent the mean of three independent experiments with five replicates each. The effect of each peptide was determined in the presence of the following concentrations of α-MSH, estimated to be the EC<sub>90</sub> value for each receptor: 10 nM for MC1R, 30 nM for MC3R, 70 nM for MC4R, and 500 nM for MC5R.
<sup>2</sup> ND; not determined.
<sup>3</sup> NC, partial activity, but not calculatable at ligand concentrations assessed
<sup>4</sup> NA; no activity at the compound concentrations assayed.

**Figure 1.**
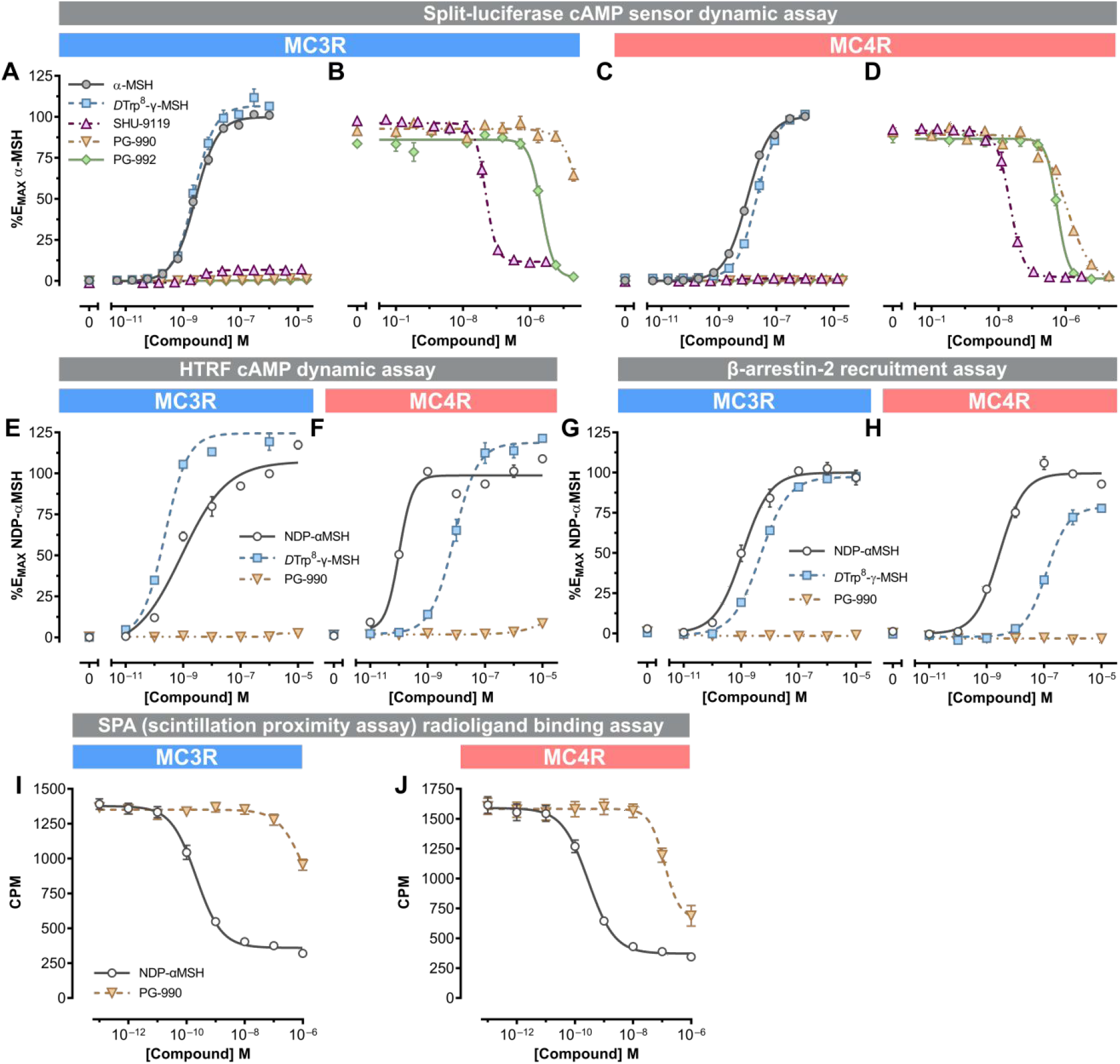
Pharmacological characterization of PG-990 and PG-992 compared to the melanocortin ligands α-MSH, NDP-α-MSH, DTrp^8^-γ-MSH, and SHU9119. The agonist (**A, C**) and antagonist (**B, D**) activities at the hMC3R (**A, B**) and hMC4R (**C, D**) were determined by a split-luciferase cAMP sensor dynamic assay in HEK-293 cells stably expressing the hMC3R or hMC4R and the GScAMP22F cAMP sensor. The antagonist activities (**B, D**) were determined in the presence of EC_80_ to EC_90_ concentration of (30 nM for hMC3R and 70 nM for hMC4R) α-MSH. Each data point represents the mean ± SEM of a representative experiment from three independent experiments with five replicates each. The EC_50_ and IC_50_ values (mean and SD) from all 3 independent experiments are found in Tables 1&2. Homogeneous time-resolved fluorescence-based dynamic cAMP assays are shown in panels **E** (hMC3R) and **F** (hMC4R). Panels showing a β-lactamase complementation assay for β-arrestin2 recruitment are depicted in **G** and **H**. Competition SPA radioligand binding assays in the presence of 80 pM of [^125^I][Nle^4^, DPhe^7^]-α-MSH are shown in panels **I** (hMC3R) and **J** (hMC4R). The data in panels **E** through **J** represent the mean ± SEM from one of two independent experiments performed in duplicate. All the data depicted in this figure were fit by a four-parameter sigmoid model.

Independent experiments, performed at Novo Nordisk in 2019 prior to knowledge of our results, reproduced these findings for PG-990. No significant hMC3R or hMC4R agonist activity was observed (Figure 1 E,F) in an assay for coupling of the receptors to G_αs_ using an antibody-based cAMP detection system (PerkinElmer, Waltham, MA) at peptide concentrations up to 10^−5^ M. Further, an orthogonal assay, based on ligand-activated receptor recruitment of β-arrestin2 (Eurofins/DiscoverX, St. Charles, MO) demonstrated no detectable agonist activity for PG-990 at either the hMC3R or hMC4R at peptide concentrations up to 10^−5^ M (Figure 1G, H). Competition binding experiments to BK cell membranes expressing the hMC3R or hMC4R, using [^125^I][Tyr^2^][Nle^4^–D-Phe^7^]-α-MSH as tracer demonstrated weak binding of PG-990 in the micromolar range (Figure 1I, J).

Analysis of PG-990 at the four human melanocortin receptors demonstrated that this peptide has weak agonist activity at hMC1R and hMC5R (Figure 2). Four independent replications of these concentration-response curves, performed at the hMC1R, hMC3R, hMC4R, and hMC5R, demonstrate the reproducibility of this assay (Supplemental Figure 1), and average EC_50_ values from the control α-MSH curves are reported in Supplemental Table 1. The activity of peptides at the hMC2R (ACTHR) is not reported in this manuscript since binding to the MC2R requires a portion of the proopiomelanocortin peptide sequence carboxyterminal to the 13 amino acid α-MSH sequence, upon which all the peptides described in the manuscript are based.

**Figure 2.**
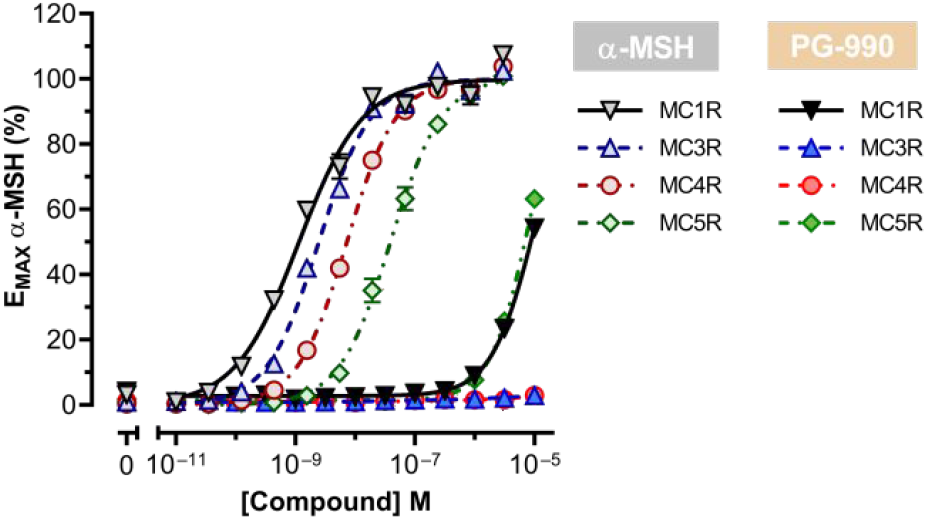
Comparison of the agonist activity as intracellular cAMP level stimulation by α-MSH and PG-990 at hMC1R, hMC3R, hMC4R, and hMC5R. HEK293 cells stably expressing the GScAMP22F split luciferase cAMP sensor (Promega) were transiently transfected with the indicated human melanocortin receptors. Each data point represents the mean ± SEM from the aggregate of four independent experiments performed in triplicate. Y-axis values are normalized to the maximum response of α-MSH for each melanocortin receptor subtype.

### PCR characterization of the University of Michigan and University of Arizona cell lines

Peptides designed and synthesized (by PG) were initially characterized pharmacologically at the University of Arizona (MC) ^25^. Thus, to determine the source of the differences between the published pharmacological results for PG-990 and PG-992 ^25^, and the results obtained herein at University of Michigan, cell lines were exchanged (between MC and RDC). HEK293 cells do not exhibit endogenous expression of any of the melanocortin receptors. Since cell lines stably transfected with expression vectors containing receptor cDNAs yield extremely high levels of receptor mRNA comparable to the levels of the actin gene, the receptor subtype expressed by each cell line can be assessed by quantitative RT-PCR. In this assay, cycle threshold (CT) values, inversely proportional to the amount of target nucleic acid, are defined as the number of PCR cycles required for the signal to exceed background levels. Even though at least two copies of each receptor sequence may be found in HEK293 genomic DNA, the thousands of copies of receptor mRNA, converted to cDNA, will be detectable at a CT value well below that required for potential detection of any contaminating genomic DNA.

Unique PCR oligonucleotide sets were synthesized based on the published and validated pairs curated by the PrimerBank database ^27^ for the hMC3R, hMC4R, hMC5R,and hMC1R (Table 4). PCR oligonucleotides for a highly expressed housekeeping gene (actin) were used to define the expected level of receptor gene expression. CT values representing amplification of endogenous receptor genomic DNA were obtained by amplification of nucleic acid from untransfected HEK293 cells using the receptor-specific oligo pairs. Multiple qPCR experiments were then performed by four different investigators, with high levels of expression represented by actin and the lowest cycle number required for signal in several experiments using untransfected cells indicated by the dashed line (Figure 3). These experiments confirmed that the hMC1R cell line from both University of Arizona and the University of Michigan expressed the hMC1R and no other hMCRs. However, these experiments also demonstrated that all three of the remaining lines from the University of Arizona were mislabeled. The line labeled MC3R expressed the hMC4R, the line labeled MC4R expressed hMC3R, and the line labeled MC5R expressed hMC4R. The University of Michigan lines all correctly expressed the receptor subtypes indicated. The extremely low hMC1R signal in the Michigan MC5R cell line (2000x lower than the hMC5R signal) is within the range of negative CT values observed across the experiment as a whole.

**Table 4.**
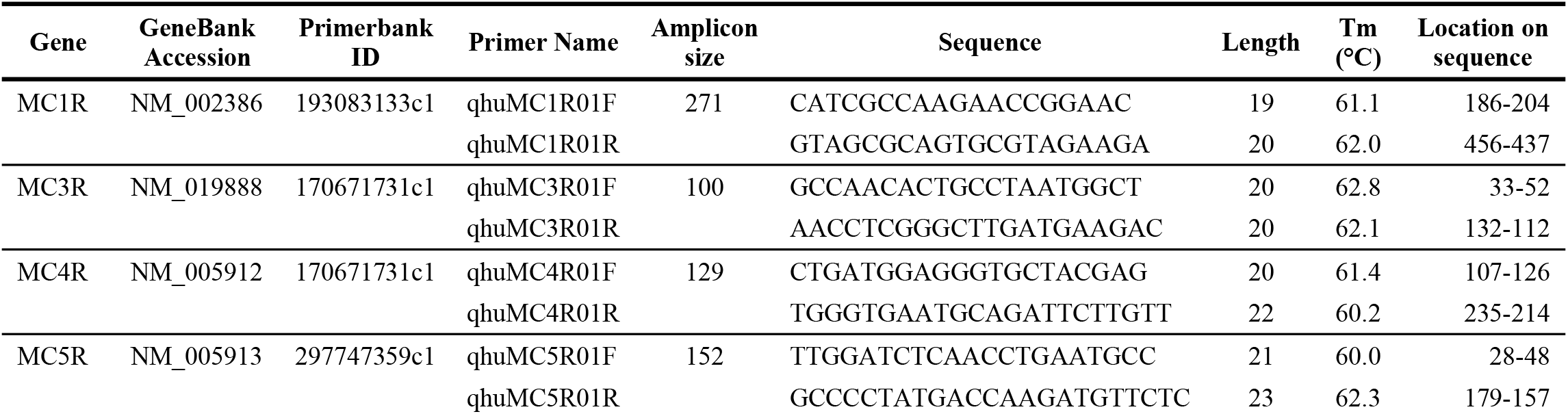
Oligonucleotide sequences used for qPCR validation of melanocortin receptor expression.

**Figure 3.**
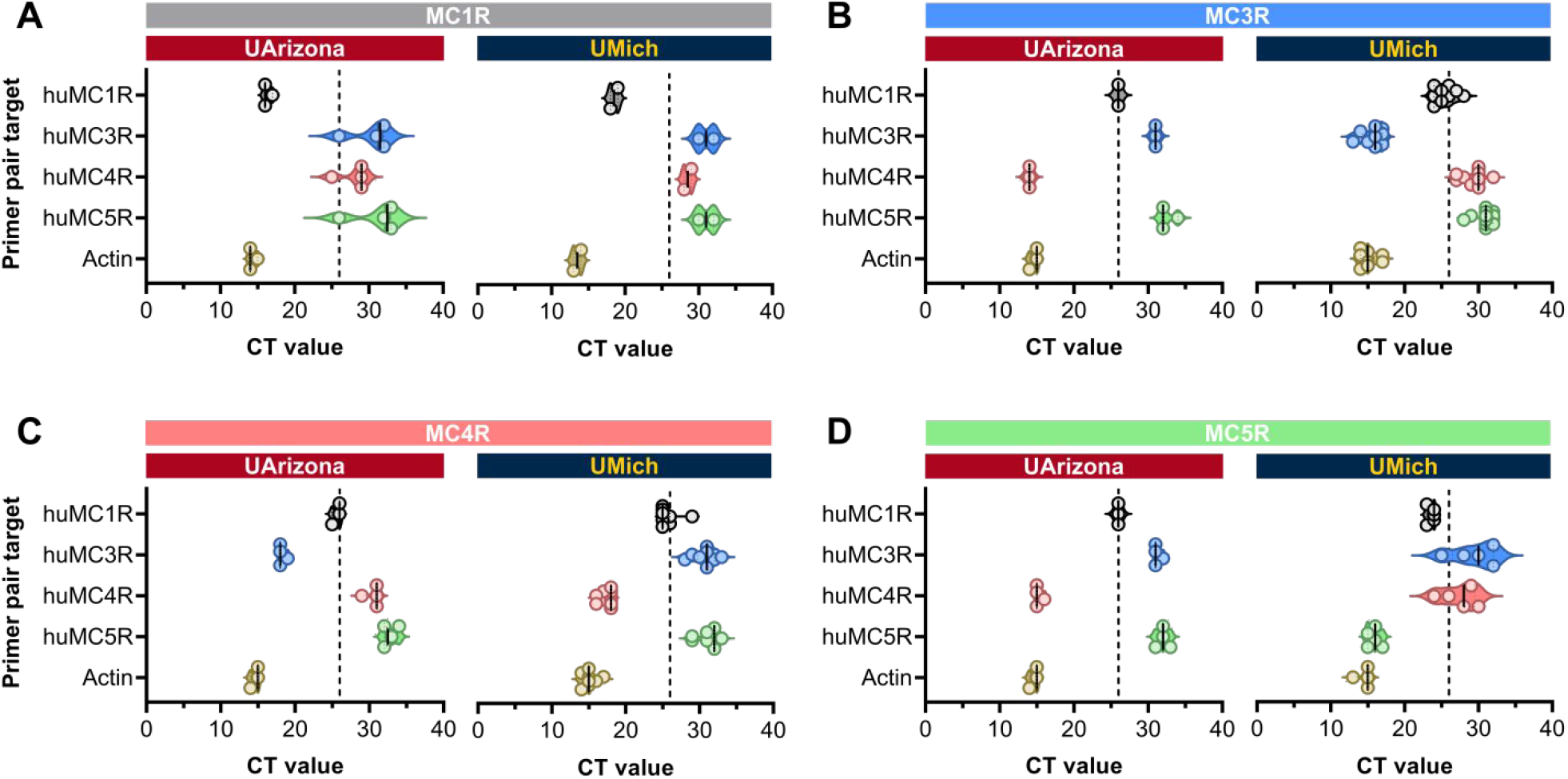
Characterization of cell lines from the University of Arizona (UArizona) and the University of Michigan (UMich) by reverse-transcriptase quantitative polymerase chain reaction experiments (qPCR). The results for cells labeled as MC1R, MC3R, MC4R, and MC5R are shown in panels **A, B, C, and D**, respectively. Each data point represents the mean CT (cycle threshold) value from an independent experiment consisting of six replicates performed two to thirteen times. The primer pairs used for each cell line are listed on the y-axis for each receptor type. The vertical dashed line on each graph indicates the lowest CT value for the receptor on the parental cell line (HEK-293, GScAMP22f) that generated the stable hMC1R, hMC3R, hMC4R, and hMC5R clones from the University of Michigan.

### Identification of potentially impacted publications

The publication reporting PG-990 and PG-992 dated back to 2015 ^25^, and thus we sought to identify additional novel peptides characterized at the University of Arizona that might require recharacterization. When initially reported, the placement of the bulky D-naphthylalanine in place of phenylalanine at position 7 (DNal(2’)^7^) of the α-MSH pharmacophore (Table 1) yielded the first potent MC4R antagonist, a cyclic heptapeptide analog of α-MSH called SHU-9119 ^28^. This widely used compound played a significant role in identifying the MC4R as a drug target for obesity ^29^, ultimately leading to the FDA-approved therapeutic, Imcivree ^30^. Interestingly, SHU-9119 remained a full agonist at the hMC1R and hMC5R and had weak partial agonist activity at the hMC3R ^28^. As this was reported in 1995, and other publications also documented the association between DNal(2’)^7^ and MC3R/MC4R antagonism or weak partial agonism ^31^, we sought to determine if the DNal(2’)^7^ residue might be a general marker of MC3R/MC4R antagonism. If so, we might then have a diagnostic tool for peptide mischaracterization over history since this DNal(2’)^7^ change was frequently included in many series of melanocortin peptides designed in a variety of labs and then characterized pharmacologically at the University of Arizona. Indeed, the DNal(2’)^7^ residue in both PG-990 and PG-992 might then explain the hMC3R and hMC4R antagonist activities seen for both peptides (Figure 1), along with the agonist activities at hMC1R and hMC5R (Figure 2). To test this hypothesis, we (TKS, Courage Therapeutics) started with three commonly used melanocortin peptide templates, a linear peptide, a 7-membered cyclic lactam, and a 7-membered cyclic disulfide. Identical (cyclic lactam and disulfide) and linear melanocortin peptides were prepared with either a DPhe or DNal(2’) at position 7, relative to the Phe position of the native α-MSH. All DPhe^7^ versions were potent agonists at hMC3R and hMC4R, while the DNal(2’)^7^ replacement uniformly produced potent hMC3R and hMC4R antagonists, with weak partial agonist curve profiles (maximum agonist activity below 20% of that observed for α-MSH) in some cases (Figure 4). The EC_50_ and IC_50_ values for all six peptides can be seen in Supplemental Table 2. As reported earlier for SHU9119 ^28^, the DNal(2’)^7^ replacement produces weak partial agonism at the MC3R and/or MC4R in some templates, which correlates with the absence of complete antagonism seen in Figure 4.

**Figure 4.**
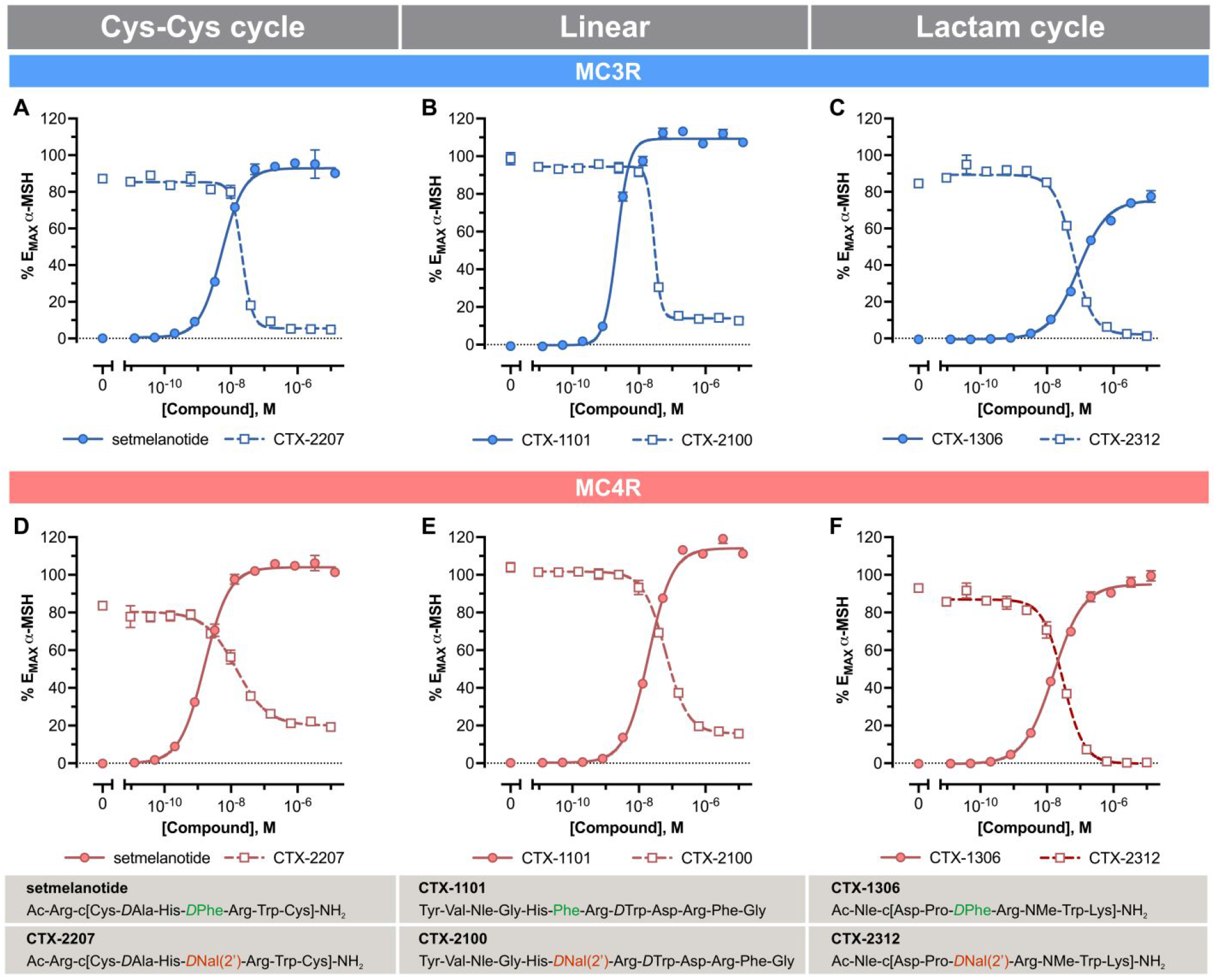
Different peptide backbones do not affect the agonist or antagonist/partial agonist activity conferred by DPhe^7^ or DNal(2’)^7^ respectively on hMC3R (**A-C**) or MC4R (**D-F**). Agonist activities determined as cAMP responses for Cys-Cys cyclic (**A**,**D**), linear (**B**,**E**), or lactam cyclic (**C**,**F**) peptides relative to the E_MAX_ for α-MSH are shown. Each data point represents the mean ± SEM of a representative experiment repeated twice with three replicates each; except for CTX-1101 which was repeated once (one experiment with three replicates) due to the limited amount of peptide available. Agonist concentration-response curves for D-Trp^8^-*γ*-MSH, a peptide related to CTX-1101, differing only by a Nle-to-Met substitution outside the tetrapeptide His-Phe-Arg-Trp pharmacophore, were repeated three times, and yielded 100% activation with similar EC_50_ values (Supplemental Figure 2).

This finding suggested that the DNal(2’)^7^ replacement may be used as a marker for peptides likely to be hMC3R/hMC4R antagonists (or very weak partial agonists), thus a tool for identifying mischaracterized peptides. Using this tool, we then screened PubMed for DNal(2’)^7^-containing peptides reported to be hMC3R or hMC4R agonists with greater than 50% agonist efficacy. From 26 papers published reporting novel melanocortin peptide structures characterized pharmacologically at Arizona ^25, 32-56^, we identified nine publications with at least 14 peptides with a DNal(2’)^7^ replacement, with the earliest paper dating back to 2002 (Supplemental Table 3). In one paper published in 2003 ^34^, we noted that two DNal(2’)^7^-containing *γ*-MSH peptide analogs, analogue **11** (Tyr-Val-Nle-Gly-His-D-Nal(2’)-Arg-Trp-Asp-Arg-Phe-Gly-NH_2_), and analogue **13** (Tyr-Val-Nle-Gly-Pro-D-Nal(2’)-Arg-Trp-Asp-Arg-Phe-Gly-NH_2_), were reported to have 100% maximal agonist activity at the hMC4R, with EC_50_ values of 24nM and 50nM, respectively, and no detectable agonist activity at the hMC5R. We show here (Figure 5A-D) that these compounds are both potent and near full agonists of the hMC5R and full antagonists of the hMC4R; no hMC4R agonist activity is detected with either peptide, while only weak partial agonist activity is detected for analog **11** at the hMC3R (<20% Emax). To confirm this finding, an independent laboratory (YXT) at Auburn University also characterized analogs **11** and **13**, using a double-blind methodology to characterize the two peptides plus α-MSH. The agonist activity of these peptides at the hMC3R, hMC4R, and hMC5R was characterized using transient expression of the receptors and a cAMP RIA detection method ^20^. The results were uncoded by a third party. As can be seen, both analogs **11** and **13** have full hMC5R agonist activity, although curiously, the cAMP endpoint assay appears to register a much lower EC_50_ value than the split luciferase assay. No agonist activity at the hMC3R or hMC4R was observed with this assay (Figure 5E-F). Thus, incorrectly reported peptide pharmacology dates back to 2003. Attribution of agonist activity of these peptides at the hMC3R and hMC4R suggests that at the time of this work written in 2003 ^34^, vials of these cells may have expressed the hMC5R.

**Figure 5.**
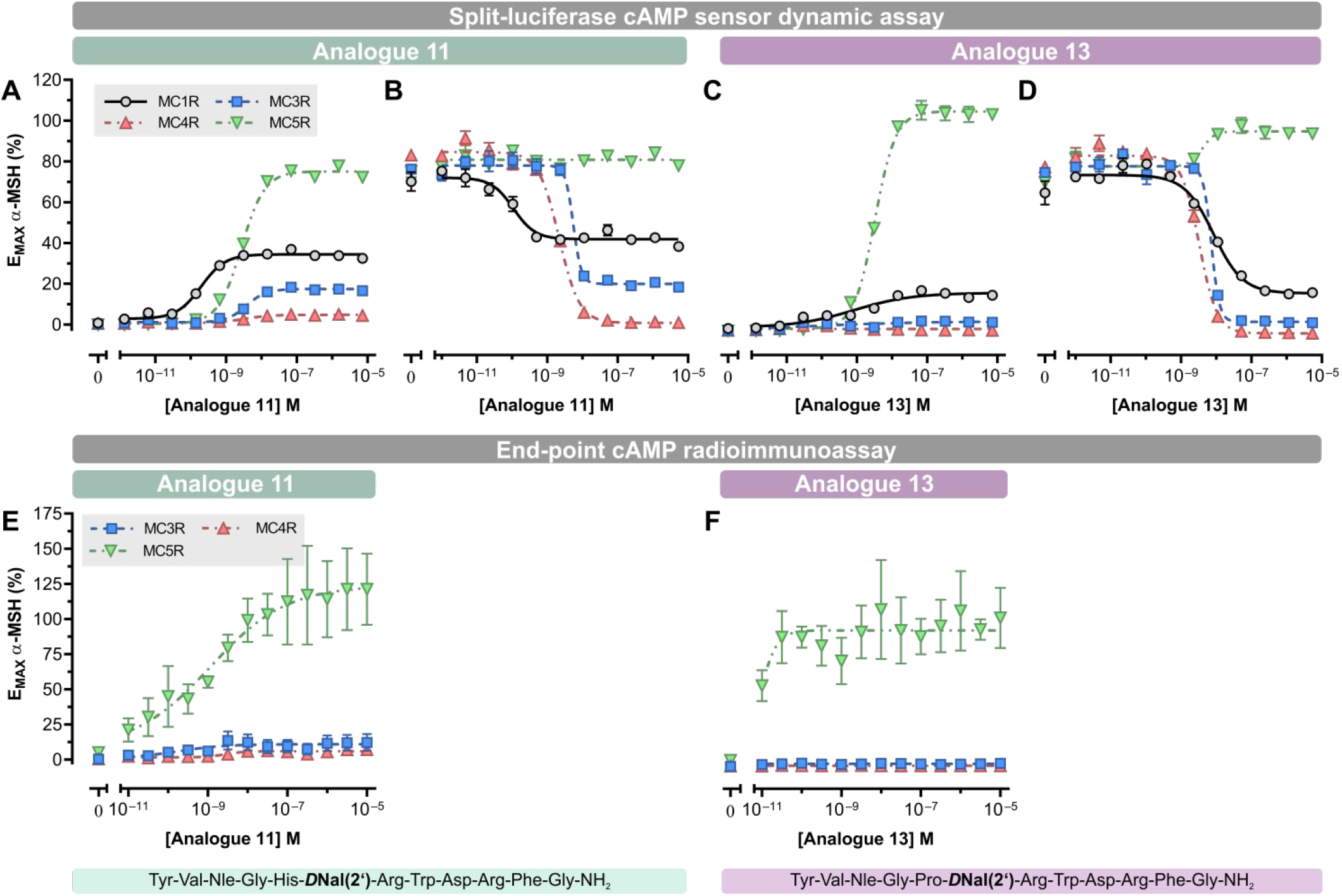
Pharmacological characterization of the indicated compounds showing agonist (**A, C**) and antagonist (**B**,**D**) activities at the hMC1-, hMC3-, hMC4-. and hMC5R using a split-luciferase cAMP sensor-based assay (**A** through **D**). The different compound cAMP responses relative to α-MSH are shown. Antagonist activity (**B**,**D**) was determined in the presence of a concentration of α-MSH equivalent to the EC_90_ for each receptor (10 nM for hMC1R, 30 nM for hMC3R, 70 nM for hMC4R, and 500 nM for hMC5R). Each data point represents the mean ± SEM of one of three independent experiments, with three replicates each. Experiments in **A** and **C**, performed at Michigan, were replicated at Auburn for hMC3R, hMC4R, and hMC5R with an end-point cAMP radioimmunoassay (RIA) -based procedure (**E**,**F**), with each data point representing the mean ± SEM of three independent experiments, with two replicates each. In these experiments using transient transfection, the data represent the mean ± SEM of one of three independent experiments performed in duplicate.

## Discussion and Conclusions

We report the systematic mischaracterization of the receptor subtype pharmacological properties of melanocortin peptides, reported in publications dating as far back as 2003 from the University of Arizona. The analysis here, using the hallmark hMC3R/hMC4R antagonist properties of DNal(2’)^7^ containing peptides, suggests that the lines, provided initially (to VJH) by one of us (RDC) in 1999 remained correctly labeled until 2002, given a report of multiple DNal(2’)^7^ peptides exhibiting hMC3R and hMC4R antagonist activity in a 2002 publication ^45^, but somehow became incorrectly labeled around that time, given the mischaracterization of peptides analogs **11** and **13** from Balse-Srinivasan et al, reported in that year ^34^, and potentially one DNal(2’)^7^ peptide (Supplemental Table 3) reported as an agonist in 2002 ^45^. The mischaracterization may not extend to the MC1R activities of peptides reported since the hMC1R expressing cell line from the University of Arizona was validated to express this receptor. The three other cell lines provided by the Arizona investigators in 2019, reported to specifically express hMC3R, hMC4R, and hMC5R, were all found to be incorrectly labeled. According to their labels, all four cell lines in use at the University of Arizona and provided (to RDC) in 2019 were at passage number 30 or greater. No obvious cross-contamination of cell lines was apparent by qPCR; thus the error may have resulted from mislabeling of cell plates or vials and the absence of an appropriate cell validation protocol. To eliminate such problems, a robust and straightforward protocol for validation of melanocortin receptor subtype expression by qPCR is provided here (see methods).

Four DNal(2’)^7^ peptides reported in two different publications ^25, 34^, dating back to 2003, are shown here to have been mischaracterized pharmacologically. The *bona fide* pharmacological properties of these peptides shown here cannot be simply explained by a single mislabeling event of cell vials provided to the University of Michigan by the University of Arizona in 2019. If this were the case, the University of Arizona would have reported PG-990 and PG-992 only to have agonist activity at MC1R since the MC3R, MC4R, and MC5R lines actually expressed MC4R, MC3R, and MC4R, respectively. Thus, the data suggest multiple mislabeling events over time.

The data here also suggest that most melanocortin peptides with a DNal(2’)^7^ residue will be hMC3R/hMC4R antagonists, with varying degrees of hMC1R and hMC5R agonist activity, and weak partial agonism, in some cases, at the hMC3R and/or hMC4R. The recent X-ray crystal structure of the inactive hMC4R bound to the DNal(2’)^7^ containing antagonist SHU9119 ^22^, and cryo-EM structure bound to a DPhe^7^-containing agonist ^57^, along with mutational data ^58^ provide a molecular explanation for the ability of DNal(2’)^7^ to antagonize hMC3R and hMC4R, but not hMC1R or hMC5R. The findings reported here suggest that the systematic pharmacological mischaracterization of the receptor subtype activity of melanocortin peptides analyzed at the University of Arizona extends to melanocortin peptide pharmacology published as far back as 2003. Investigators should thus recharacterize any peptides of interest from these publications before conducting any further research with them.

## Experimental Section

Dr. Paolo Grieco kindly provided PG-990 and PG-992. Setmelanotide, DTrp^8^-*γ*-MSH, CTX-1101, CTX-1306, CTX-2207, CTX2100, and CTX-2312 were provided by Courage Therapeutics, Inc., after synthesis by Vivitide (Gardner, MA). Peptide analog **11**, and analog **13** ^34^ were obtained from Vivitide. Peptide PG-990, characterized by scientists from Novo Nordisk, was prepared in two batches at Novo Nordisk. All peptides in this study were >95% pure by analytical RP-HPLC and had a mass within 1% of the calculated weight, as determined by mass spectrometry.

### Pharmacological Assays

#### Determination of intracellular cAMP levels in live cells (Univ. Michigan)

The methodology for determining cAMP levels in live cells is described in detail elsewhere ^22^. In brief, a cAMP split-luciferase reporter (GScAMP22F) stably-expressing cell line (Promega, Madison, WI) was transfected with hMC1R, hMC3R, hMC4R, and hMC5R expression vectors using lipofectamine, and stable clonal cell lines were selected for use in this study. The plasmids used for transient transfections were obtained from the cDNA Resource Center (www.cdna.org). To determine cAMP levels, cells were seeded at a density of 20,000 cells per well using 384-well poly-D lysine-coated, clear bottom, and black-wall assay plates (Corning Inc. Corning, NJ). Cells were allowed to attach to the plates for 18 to 24 h, after which growth media was removed and 20 µl of 4% D-luciferin (Promega) in CO_2_ independent medium (Thermo Fisher Scientific) was added to each well. The luciferase substrate was allowed to permeate the cells for 120 min at 37 °C. Intracellular cAMP levels were measured using an FDSS 7000EX Functional Drug Screening System (Hamamatsu Photonics, Hamamatsu, Japan).

#### Determination of intracellular cAMP levels by cAMP RIA (Auburn University)

Human embryonic kidney (HEK) 293T cells (ATCC, Manassas, VA, USA) were cultured at 37 °C in a 5% CO_2_-humidified atmosphere. Cells were transiently transfected with hMC3R, hMC4R, or hMC5R (0.25 mg/mL) using a calcium phosphate precipitation method. The final concentration of experimental peptides and α-MSH used were 10 pM to 10 µM. cAMP signaling assay was performed following cell lysis by radioimmunoassay as described previously ^20^. Data are mean ± S.E.M. from three separate experiments, with duplicate measurements within each experiment.

#### Determination of intracellular cAMP levels using an antibody-based FRET method (Novo Nordisk)

The assays were performed in 96-well white opaque plates. Compounds and cells were diluted in buffer (DMEM w/o phenol red, 10 mM Hepes, 1x Glutamine, 0.1% (w/v) ovalbumin, 1mM IBMX). 25 µl of appropriate dilutions of test compounds were added in the respective wells. Compounds were tested in duplicates in each experiment. The assay was initiated by adding 25 µl suspension (4000 cells/well) of BHK (baby hamster kidney) cells stably expressing the human MC3 or MC4 receptor and incubated for 30 minutes at 25°C. cAMP induction was subsequently measured by cAMP Gs dynamic HTRF kit from CisBio according to the protocol provided by the vendor. The plates were read on Mithras LB 940 provided by Berthold Technologies.

#### DiscoverX β-Arrestin recruitment assay procedure (Novo Nordisk)

The following kits were purchased from Eurofins/DiscoverX: PathHunter® eXpress MC3R U2OS β-Arrestin GPCR Assay (#93-0984E3) PathHunter® eXpress MC4R U2OS β-Arrestin GPCR Assay (#93-0211E3). Suitable dilutions of test compounds were tested in duplicates in each experiment according to the protocol provided by the vendor. *SPA binding assay procedure:* Membranes were prepared from BHK cell lines stably expressing human MC3R or MC4R. Cell pellets were homogenised in ice cold buffer (20 mM Hepes, 5 mM MgCI_2_, 1 mg/ml Bacitracin, pH 7.1 and one cOmplete™ Protease Inhibitor Cocktail Tablet (Roche Applied Science) per 25 ml and centrifuged at 25000 g at 4°C for 10 minutes. The supernatant was discarded, and the pellets were re-suspended in the buffer, and then homogenised and centrifuged two more times. The pellets were pooled, and the final pellet was re-suspended in buffer, aliquoted and subsequently stored at −80°C.

The SPA Binding assays were performed in 96-well white opaque plates. Each well contained: 0.5 mg PVT-WGA SPA beads, hMC3 or hMC4 receptor expressing membrane diluted to give approximately 10% specific tracer binding, 50000 dpm [125I][Tyr^2^][Nle^4^–D-Phe^7^]-α-MSH and relevant dilutions of test compounds. Compounds were tested in duplicates in each experiment. The final volume in each well was 200 µl. The assay buffer was 25 mM Hepes, pH 7.0, containing 1.5 mM CaCl_2_, 1 mM MgSO_4_, 0.21% (w/v) ovalbumin, 1 mM 1,10-Phenanthroline and one cOmplete™ Protease Inhibitor Cocktail Tablet per 100 ml. Plates were incubated overnight (22-24 hours) at room temperature before counting for 2 minutes per well, using a TopCount NXT scintillation counter.

### qPCR assays

HEK293 cells stably expressing the human melanocortin receptors (hMC1R, hMC3R, hMC4R, and hMC5R) from the University of Arizona and the University of Michigan were grown in Dulbecco’s modified Eagle’s medium (DMEM) supplemented with 10% FBS, and hygromycin B and/or Geneticin for selection. Cells were maintained at 37 °C in a humidified incubator in the presence of 5% CO_2_. Total RNA was isolated from the cell lines using TRIzol® RNA isolation reagent (Invitrogen, Carlsbad CA) according to the manufacturer’s instructions. The total RNA concentration was determined by spectrophotometry at a peak absorbance of 260 nm, and quality was assessed by obtaining the absorbance ratio between the 280 and 260 nm absorbance values. cDNA was synthesized from 1 *μ*g of total RNA using a High Capacity cDNA reverse transcription kit (Applied Biosystems, Foster City, CA) in a final volume of 20 μL. The reaction was incubated at 25 °C for 10 min, 37 °C for 120 min, 85 °C for 5 min, and kept at −20 °C until ready for use. All samples were diluted by 1:10 with DNase and RNase-free distilled water to obtain the required concentration for RT-qPCR analysis. The primers targeting the human melanocortin receptors (MC1R, MC3R, MC4R, and MC5R), and targeting a housekeeping gene expressed at high levels (actin) were obtained from the PrimerBank database ^27^. Table 4 summarizes the sequences and other parameters for the primers used in this study. Real-time semi-quantitative PCR - (RT-qPCR) was performed using an Applied Biosystems QuantStudio 5 Real-Time PCR System. Each 10 μL reaction contained 5 μL of PowerSYBR Green PCR Master Mix (Applied Biosystems), forward and reverse primers at a final concentration of 0.2 μM, and 2 μL of the specific cDNA for each cell sample. The RT-qPCR amplification program consisted of a 10-min pre-denaturation step at 95 °C, 40 cycles of denaturation at 95 °C for 15 sec, and annealing/extension at 60 °C for 1 min. As the sole purpose of these experiments was to demonstrate the predominant receptor type expressed by each cell line, we compiled the raw cycle threshold (CT) values obtained at different times from several rounds of experimentation by different investigators.

### Literature Analyses

A list of all previous publications, including Drs. Minying Cai and Victor J. Hruby as authors was compiled from the PubMed database using the joint author terms for the search. This list was refined by inspection of each article to include publications presenting new compound pharmacological characterizations. Review articles were excluded from the list. Compounds in Supplemental Table 3 were extracted from the entire collection of 346 published peptides in these 26 publications based on the following criterion: the compound(s) possessed DNal(2’) at position 7 in the MSH amino acid sequences and were found to have agonist activity (≥50% E_MAX_) at the MC3R or MC4R. Compounds that met this criterion were then compiled into a separate Microsoft Excel spreadsheet and organized into reverse chronological order based on publication year. Supplemental Table 3 includes 1 to 3 compound(s) from each article possessing DNal(2’) at position 7 of the α-MSH amino acid sequence and the following compound information: peptide number; peptide name; peptide sequence; and notes on the found activity of the peptide.

## Acknowledgments

This work was funded in part by NIH DK126715 (RDC), NIHR41124449 (TS), NIH R41DK127817 (TS), Courage Therapeutics (RDC and TS), and the Klarman Foundation (RDC). We thank Drs. Victor J. Hruby and Minying Cai for providing cell lines from their labs. We thank Dr. Birgitte S. Wulff (Novo Nordisk A/S) for providing data and discussion relevant to this research.

## Special note from the authors

Professor Hruby and his team members have made many outstanding contributions to melanocortin receptor-targeted peptide design and the development of melanocortin drugs over the past 40 years. Contributions cited in this paper include the design of the first super-potent melanocortin peptide agonists and the first MC3R/MC4R antagonist. These basic principles of melanocortin peptide design continue to guide the field today. This manuscript is not meant to detract from this laudatory body of scientific work. Instead, we report only errors in characterizing receptor subtype specificity properties for some peptides with melanocortin activity published by this group.

### ABBREVIATIONS USED

α-MSH: alpha melanocyte stimulating hormone
CT: cycle threshold
DNal: D-Naphthylalanine
GαS: stimulatory G protein alpha subunit
hMC1R: human melanocortin-1 receptor
hMC2R: human melanocortin-2 receptor
hMC3R: human melanocortin-3 receptor
hMC4R: human melanocortin-4 receptor
hMC5R: human melanocortin-5 receptor
qPCR: quantitative polymerase chain reaction

## Ancillary Information

Supporting Information: Individual pharmacological experiments and EC_50_ values validating reproducibility of the findings, individual EC_50_ and IC_50_ values for key experimental peptides, Table of D-Nal(2’)7 containing peptides reported to have MC3R or MC4R agonist activity and their papers of origin, published papers, and quality control data for peptides used in this study.

Figure S1

Figure S2

Table S1

Table S2

Table S3

Table S4

Peptide QC Data

## TABLE OF CONTENTS GRAPHIC

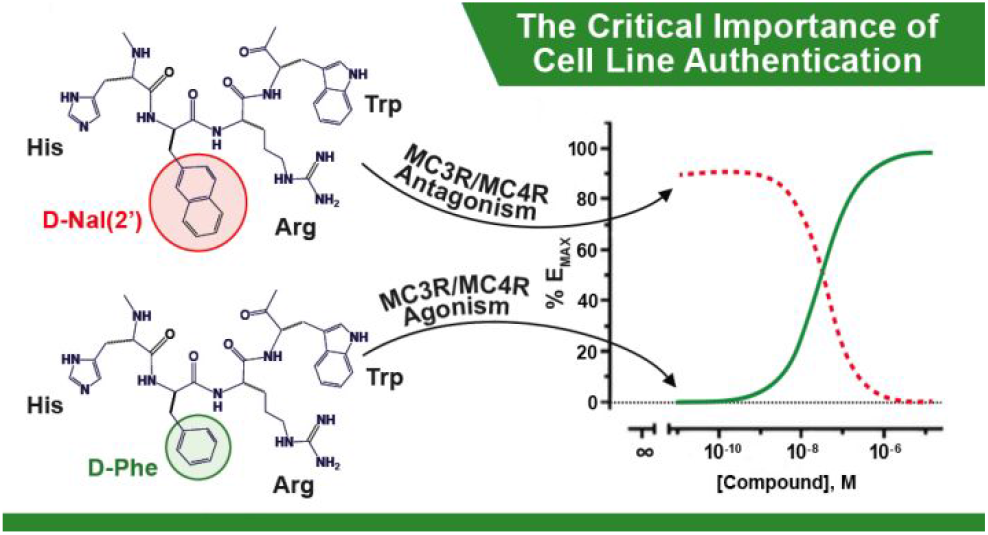

