## Supplemental Files for "Demonstration of a common DPhe^7^ to DNal(2’)^7^ peptide ligand antagonist switch for the melanocortin-3 and melanocortin-4 receptors identifies systematic mischaracterization of the pharmacological properties of melanocortin peptides"

##### Contents

Figure S1
Figure S2
Table S1
Table S2
Table S3
Peptide QC Data

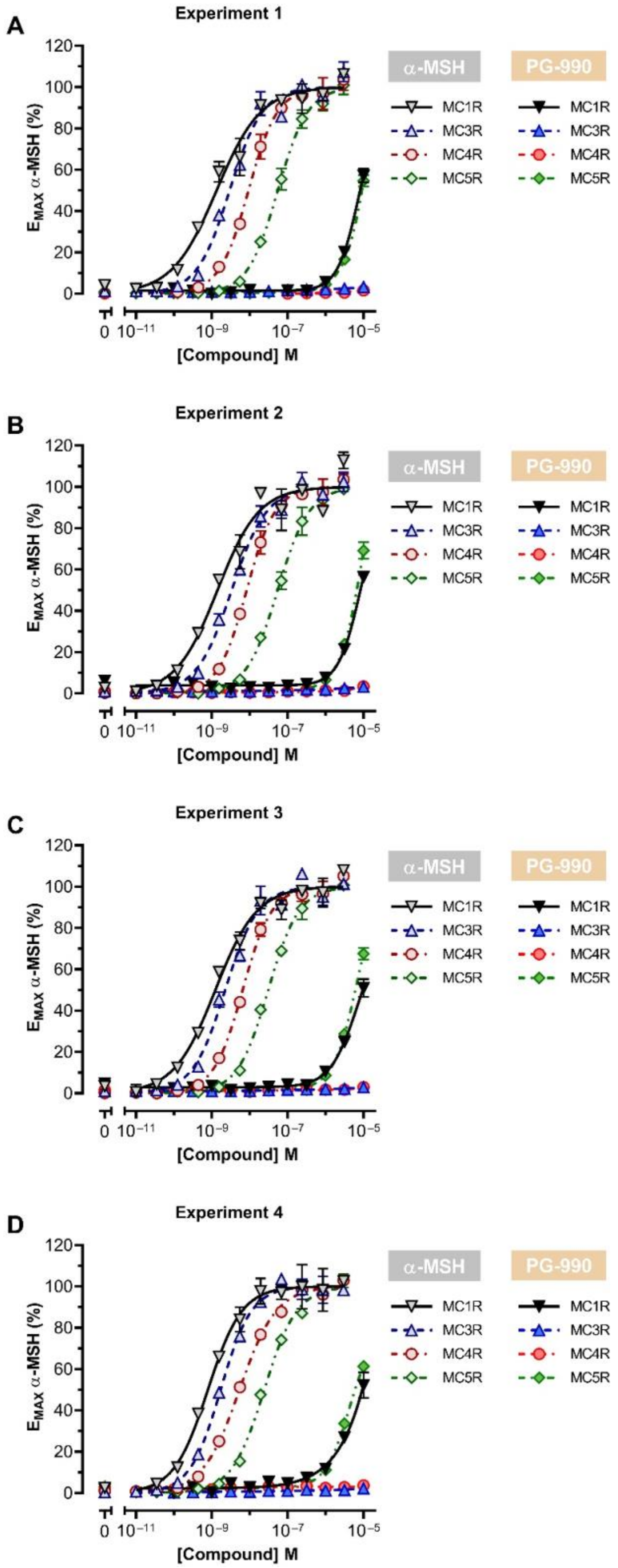


**Figure S1:** Related to **Figure 2**. Individual experiments in A, B, C, and D averaged to generate the graph in Figure 2. Each data point represents the mean ± SEM of an individual experiment performed in triplicate. Y-axis values are normalized to the maximum response of α-MSH for each melanocortin receptor subtype.

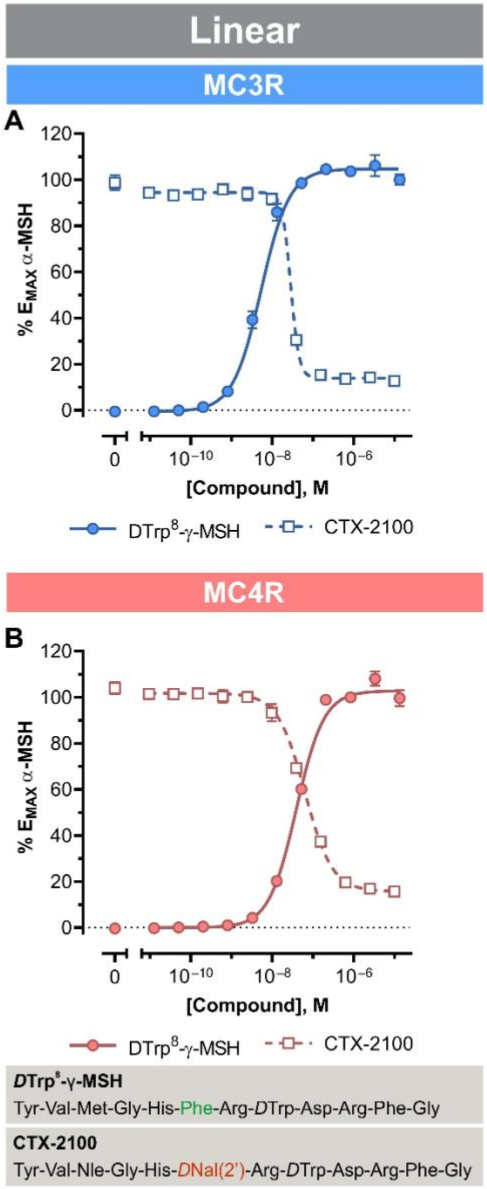


**Figure S2:** Related to **Figure 3**. Different peptide backbones do not affect the agonist or antagonist/ partial agonist activity conferred by DPhe or DNal(2') respectively on MC3- (**A**) or MC4R (**B**). Each data point represents the mean ± SEM of a representative experiment repeated twice with three replicates each.

| Receptor type | $pEC_{50}$ Exp. 1[3] | $pEC_{50}$ Exp. 2 | $pEC_{50}$ Exp. 3 | $pEC_{50}$ Exp. 4 | Mean $pEC_{50}$ ± SEM ($EC_{50}$, nM)[4] |
|---|---|---|---|---|---|
| **MC1R** | 8.873 | 8.850 | 8.905 | 9.103 | 8.933 ± 0.058 (***1.2***) |
| **MC3R** | 8.530 | 8.476 | 8.646 | 8.766 | 8.605 ± 0.064 (***2.5***) |
| **MC4R** | 8.032 | 8.080 | 8.186 | 8.270 | 8.142 ± 0.053 (***7.2***) |
| **MC5R** | 7.259 | 7.263 | 7.511 | 7.617 | 7.413 ± 0.090 (***38.7***) |

Table **S1**[1]: **αMSH $EC_{50}$ values for hMC1R, hMC3R, hMC4R, and hMC5R**[2]

[1] This table is associated with Figure 2.
[2] HEK-293 cells stably expressing the GScAMP22f split-luciferase cAMP sensor were transfected with plasmids (pcDNA3.1(+) backbone) bearing the sequences of the human MC1R (RefSeq ID: NM_002386.3), MC3R (RefSeq ID: NM_019888.3), MC4R (RefSeq ID: NM_005912.2), or MC5R (RefSeq ID: NM_005913.2).
[3] Each individual experiment was performed in triplicate wells.
[4] The $pEC_{50}$ values were obtained by fitting a four-parameter sigmoid function to concentration-response curves for each specific condition as indicated.

| MC3R | | | | | |
|---|---|---|---|---|---|
| **Compound** | **$pEC_{50}$ or $pIC_{50}$ Exp. 1**[3] | **$pEC_{50}$ or $pIC_{50}$ Exp. 2**[3] | **$pEC_{50}$ or $pIC_{50}$ Exp. 3**[3] | **$pEC_{50}$ or $pIC_{50}$ Exp. 4**[3] | **Mean $pEC_{50}$ ± SEM or $pIC_{50}$ ± SEM ($EC_{50}$ or $IC_{50}$, nM)**[4] |
| **setmelanotide** | 8.258 | 8.615 | 8.583 | 8.659 | 8.529 ± 0.092 (***3.0***) |
| **CTX-2207** | 7.652 | 7.671 | - | - | 7.662 ± 0.010 (***21.8***) |
| ***D*Trp[8]-γ-MSH** | 8.350 | 8.730 | 8.940 | 8.660 | 8.670 ± 0.122 (***2.1***) |
| **CTX-2100** | 7.139 | 7.589 | - | - | 7.364 ± 0.225 (***43.3***) |
| **CTX-1306** | 7.002 | 8.209 | - | - | 7.606 ± 0.604 (***24.8***) |
| **CTX-2312** | 7.200 | 7.203 | - | - | 7.202 ± 0.002 (***62.9***) |

| MC4R | | | | | |
|---|---|---|---|---|---|
| **Compound** | **$pEC_{50}$ or $pIC_{50}$ Exp. 1**[3] | **$pEC_{50}$ or $pIC_{50}$ Exp. 2**[3] | **$pEC_{50}$ or $pIC_{50}$ Exp. 3**[3] | **$pEC_{50}$ or $pIC_{50}$ Exp. 4**[3] | **Mean $pEC_{50}$ ± SEM or $pIC_{50}$ ± SEM ($EC_{50}$ or $IC_{50}$, nM)**[4] |
| **setmelanotide** | 8.779 | 9.259 | 9.273 | 9.324 | 9.159 ± 0.127 (***0.7***) |
| **CTX-2207**[5] | 7.850 | 7.847 | | | 7.849 ± 0.001 (***14.2***) |
| ***D*Trp[8]-γ-MSH** | 7.360 | 7.345 | 7.108 | 7.272 | 7.271 ± 0.058 (***53.5***) |
| **CTX-2100**[5] | 7.220 | 7.196 | | | 7.208 ± 0.012 (***61.9***) |
| **CTX-1306** | 7.764 | 8.301 | 7.802 | | 7.956 ± 0.173 (***11.1***) |
| **CTX-2312**[5] | 7.509 | 7.530 | | | 7.520 ± 0.011 (***30.2***) |

**Table S2**[1]: **Agonist or antagonist potencies for compounds shown in Figure 4.**[2]

[1] This table is associated with Figure 3.
[2] HEK-293 cells stably expressing the GScAMP22f split-luciferase cAMP sensor and either the MC3R (RefSeq ID: NM_019888.3) or MC4R (RefSeq ID: NM_005912.2) were used.
[3] Each experiment was performed in triplicate, and two to four independent experiments were performed as indicated.
[4] The $pEC_{50}$ or $pIC_{50}$ values were obtained by fitting a four-parameter sigmoid function to concentration-response curves for each specific condition as indicated.
[5] Inhibition profiles were determined for the compounds with antagonist activity (i.e. CTX-2207, CTX-2100, and CTX-2312) after a 12 min incubation with αMSH at an EC90-equicvalent concentration adjusted for each receptor type (i.e. 30 nM for hMC3R and 70 nM for hMC4R respectively).

| Peptide #[1] | | Sequence | Notes |
|---|---|---|---|
| **Development of Macrocyclic Peptidomimetics Containing Constrained α,α-Dialkylated Amino Acids with potent & selective Activity at Human Melanocortin Receptors.**<br>Merlino et al., J Med Chem. 61(9), 4263-4269, 2018 | | | |
| 13 | pep 13 | [Aib-His-D-Nal(2')-Arg-Trp-Glu]-NH2 | **hMC5R antagonist: potent; hMC1R & hMC4R agonist: full; hMC3R agonist: partial** |
| **Discovery of Novel potent & selective Agonists at the Melanocortin-3 Receptor.**<br>Carotenuto et al., J Med Chem. 58(24), 9773-9778, 2015 | | | |
| 2 | PG-990 | Ac-Nle-c[Asp-Pro-Pro-D-Nal(2')-Arg-Trp-Lys]-NH2 | **hMC3R agonist: potent & selective** |
| 4 | PG-992 | Ac-Nle-c[Asp-Trp-Pro-D-Nal(2')-Arg-Trp-Lys]-NH2 | **hMC3R agonist: potent & selective** |
| **Systematic Backbone Conformational Constraints on a Cyclic Melanotropin Ligand Leads to Highly Selective Ligands for Multiple Melanocortin Receptors.**<br>Cai et al., J Med Chem. 58(16), 6359-6367, 2015 | | | |
| 14 | pep 14 | Ac-Nle-c[Asp-His-**D-Nal(2')**-**Arg**-Trp-Lys]-NH2 | **hMC4R agonist and hMC3R agonist: (73% activation and 78% activation, respectively)** |
| 17 | pep 17 | Ac-Nle-c[Asp-His-D-Nal(2')-**Arg**-**Trp**-**Lys**]-NH2 | **hMC3R antagonist: selective; hMC4R agonist: (67% activation)** |
| **Substitution of arginine with proline and proline derivatives in melanocyte-stimulating hormones leads to selectivity for human melanocortin 4 receptor.**<br>Qu et al., J Med Chem. 52(12), 3627-3635, 2009 | | | |
| 6 | pep 6 | Ac-Nle-c[Asp-His-D-Nal(2')-Pro-Trp-Lys]-NH2 | **hMC4R antagonist: selective (55% activation)** |
| 7 | pep 7 | Ac-Nle-c[Asp-His-D-Nal(2')-trans-Xaa-Trp-Lys]-NH2 | **hMC4R antagonist: selective; hMC3R agonist: (56% activation)** |
| **Effects of macrocycle size and rigidity on melanocortin receptor-1 and -5 selectivity in cyclic lactam alpha-melanocyte-stimulating hormone analogs.**<br>Mayorov et al., Chem Biol Drug Des. 67(5), 329-335, 2006 | | | |
| 16 | Analog 16 | Ac-c[Glu-His-D-Nal(2')-Arg-Trp-Lys]-NH2 | **hMC3R agonist: (75% activation)** |
| **Development of cyclic gamma-MSH analogues with selective hMC3R agonist and hMC3R/hMC5R antagonist activities.**<br>Mayorov et al., J Med Chem. 49(6), 1946-1952, 2006 | | | |
| 11 | Analog 11 | c[Nle-Gln-D-Nal(2')-Arg-Trp-Glu]-NH2 | **hMC4R agonist: (50% activation)** |
| **Structure-activity studies of new melanocortin peptides containing an aromatic amino acid at the N-terminal position.**<br>Grieco et al., Peptides. 27(2), 472-481, 2006 | | | |
| 8 | PG-978 | H-D-Nal-c[Asp-Pro-D-Nal(2')-Arg-Trp-Gly-Lys]-NH2 | **hMC3R antagonist and hMC4R antagonist: (85% activation @ hMC3R)** |
| **Structure-activity relationships of gamma-MSH analogues at the human melanocortin MC3, MC4, and MC5 receptors. Discovery of highly selective hMC3R, hMC4R, and hMC5R analogues.**<br>Balse-Srinivasan et al., J Med Chem. 46(23), 4965-4973, 2003 | | | |
| 11 | analog 11 | H-Tyr-Val-Nle-Gly-His-D-Nal(2')-Arg-Trp-Asp-Arg-Phe-Gly-NH2 | **hMC4R agonist: potent & selective (100% activation)** |
| 13 | analog 13 | H-Tyr-Val-Nle-Gly-Pro-D-Nal(2')-Arg-Trp-Asp-Arg-Phe-Gly-NH2 | **hMC4R agonist: potent & selective (100% activation); hMC3R agonist: (70% activation)** |
| 14 | analog 14 | H-Tyr-Val-Nle-Gly-His-D-Nal(2')-Arg-D-Nal(2')-Asp-Arg-Phe-Gly-NH2 | **hMC4R agonist: potent & selective (100% activation)** |
| **Novel cyclic templates of alpha-MSH give potent & highly selective antagonists/agonists for human melanocortin-3/4 receptors.**<br>Kavarana et al., J Med Chem. 45(12), 2644-2650, 2002 | | | |
| 5 | MK-5 | (O)C-(CH2)2-C(O)-c[His-D-Nal(2')-Arg-Trp-Lys]-NH2 | **hMC3R agonist: potent & selective (88% activation)** |

**Table S3[1]: D-Nal(2')$^{7}$-containing peptides reported as hMC3R and/or hMC4R agonists (greater than or equal to 50% Emax activity at either hMC3R or hMC4R)**

[1] peptide number references the numerical identifier used for each peptide in the original publications referenced here.

**QUALITY CONTROL SECTION**

| Peptide | Purity (%) | Calculated Mass (g/mol) | Measured Mass (g/mol) |
|---|---|---|---|
| **DTrp$^{8}$-γ-MSH** | 97.25 | 1570.8 | 1570.7 |
| **Setmelanotide** | 98.47 | 1117.3 | 1117.68 |
| **SHU-9119** | 100 | 1074.1 | 1074.55 |
| **PG-990 (PG)** | >99 | 1131.6104 | 1131.6068 |
| **PG-992 (PG)** | >99 | 1220.6369 | 1220.6338 |
| **PG-990 (Novo1A)** | 99.72 | 1131.6025 | 1131.6076 |
| **PG-990 (Novo3A M+2/2)** | 100 | 566.3013 | 566.5 |
| **CTX-2207** | 96.69 | 1167.3 | 1167.59 |
| **CTX-1101** | 97.6 | 1552.84 | 1552.7 |
| **CTX-2100** | 100 | 1602.6 | 1602.81 |
| **CTX-1306** | 99.74 | 998 | 998.57 |
| **CTX-2312** | 100 | 1048.1 | 1048.59 |
| **Analogue 11** | 99.22 | 1601.6 | 1601.77 |
| **Analogue 13** | 100 | 1561.6 | 1561.74 |

**Table S4: Quality control data for experimental peptides in this study.**

| ID | $t_R$[a] (min) | % isolated purity[a] | HRMS[b] *m/z* | |
|---|---|---|---|---|
| | | | calcd | obsd |
| **PG-990** | 12.7 | >99 | 1131.6104 | 1131.6068 |
| **PG-992** | 15.0 | >99 | 1220.6369 | 1220.6338 |

**Table S5. Analytical data of synthesized compounds PG-990 and PG-992 (from PG).**

[a]Compounds were analyzed by analytical UHPLC (Shimadzu Nexera Liquid Chromatograph LC-30AD), performed on a C18-bonded Kinetex reverse-phase column from Phenomenex (150 mm × 4.6 mm, 5 µm, 100 Å) with a flow rate of 1 mL/min and using linear gradients of MeCN (0.1% TFA) in water (0.1% TFA), from 10 to 90% over 15 min. [b]HRMS calculated and observed for $[M+H]^+$ by LTQ Orbitrap.

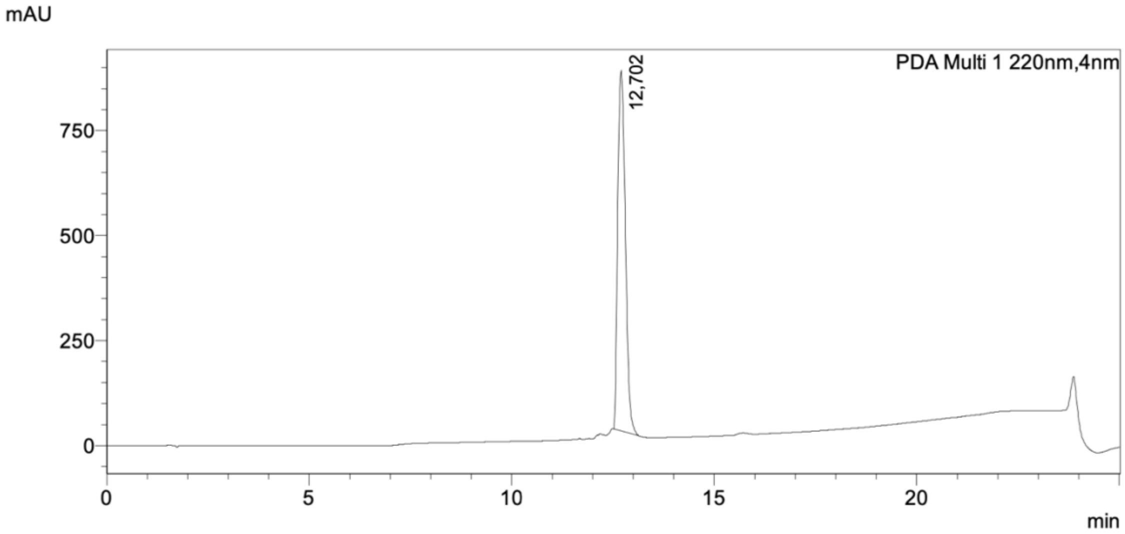


Chromatogram of **PG-990**.

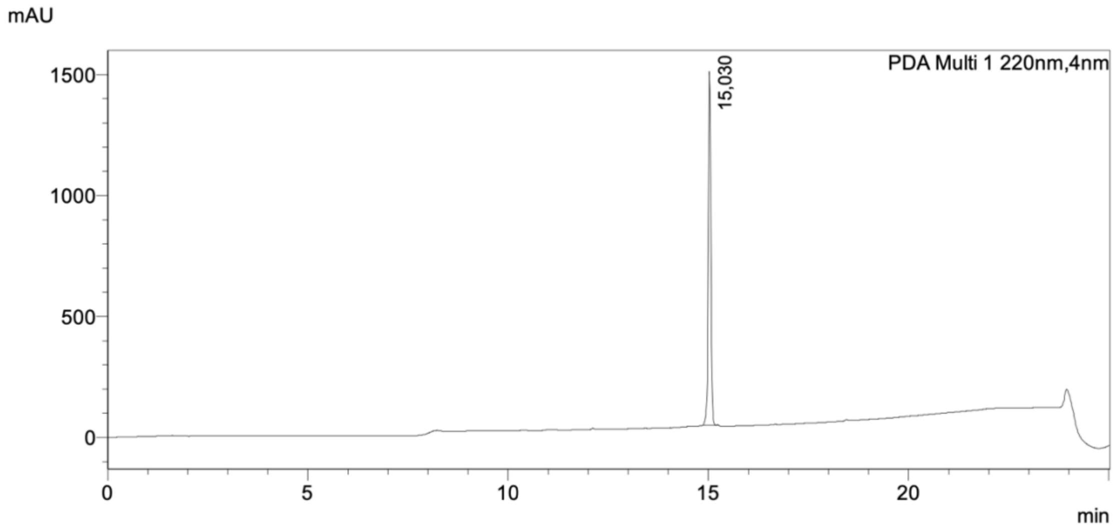


Chromatogram of **PG-992**.

### QC data for PG990 peptide used by Novo Nordisk

PG990 (in house batch 3a):
LS-MS data: observed 1131.6076, calc M+1 1131.6025 (Waters 6200 series TOF/6500 series Q-TOF B.05.00 (B5042.2))

PG990 (in house batch 1a):
LS-MS data: observed 566.5, calc (M+2)/2 566.3013 (Waters Micromass ZQ detector)

BATCH 1A

Page: 1 of 1
Project Name: Apigenex Peptidy 2

**Sample information**

**UPLC2**

**Sample: A170425/1-6 0070-0000-1350 h.p.**

Channel Description ACQUITY TUV ChA 214nm

Vial : 1:A,6 Vol. : 0,20 ul

Date Acquired 12.5.2017 10:46:59 CEST

Date Processed 12.5.2017 10:59:00 CEST

Acq Method Set :
Gr_5_60_16min_40C_0_4_K1_met_s

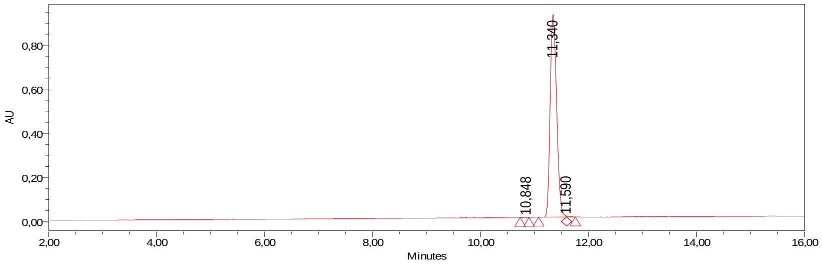


| | RT | Area | Height (µV) | % Area |
|---|---|---|---|---|
| 1 | 11,340 | 7631456 | 919373 | 99,72 |
| 2 | 11,590 | 16276 | 4286 | 0,21 |

A: 0.05% TFA in water
B: 0.05% TFA in acetonitrile
Gradient : 5--> 60% B, 16min, 0.4ml/min
Acquity UPLC BEH130, 1.7um, 2.1 x 150 mm column
column own temp. = 40 °C

BATCH 3A



#### Chromatogram and Results

##### Injection Details

| | | | |
|---|---|---|---|
| *Injection Name:* | **hst20539-706-I** | *Run Time (min):* | **15.00** |
| *Vial Number:* | **R:A2** | *Injection Volume:* | **5.00** |
| *Injection Type:* | **Unknown** | *Channel:* | **CAD_1** |
| *Calibration Level:* | | *Wavelength:* | |
| *Instrument Method:* | **CAD V10** | *Bandwidth:* | **4** |
| *Processing Method:* | **CAD V10** | *Dilution Factor:* | **1.0000** |
| *Injection Date/Time:* | **29/May/18 10:29** | *Sample Weight:* | **1.0000** |

##### Chromatogram

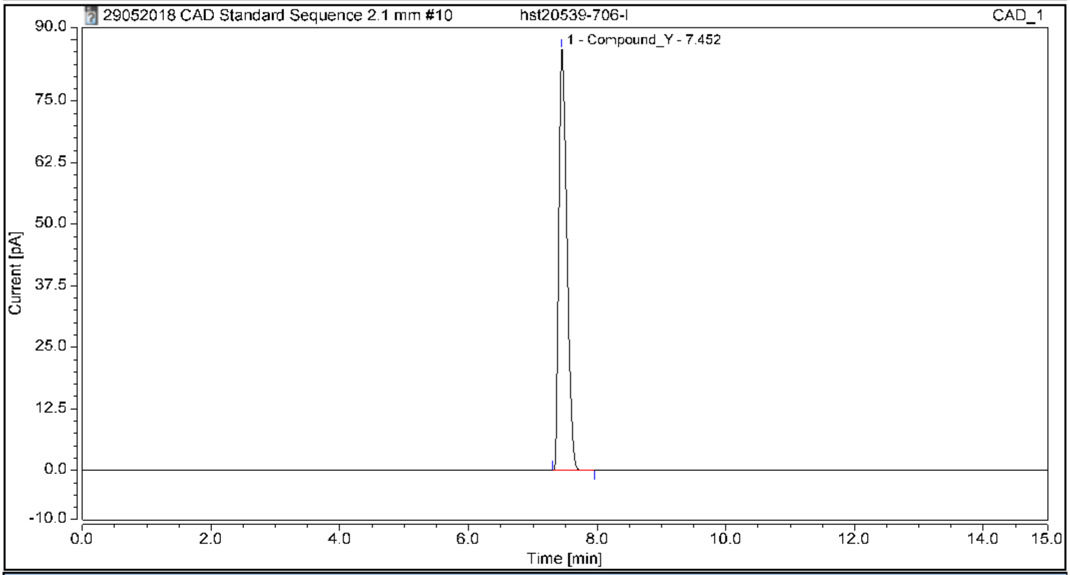


##### Integration Results

| No. | Peak Name | Retention Time min | Area pA*min | Height pA | Amount µg | Concentration mg/mL | Evaluation |
|---|---|---|---|---|---|---|---|
| n.a. | Compound_X | n.a. | n.a. | n.a. | n.a. | n.a. | n.a. |
| 1 | Compound_Y | 7.452 | 12.432 | 85.471 | 12.4095 | 2.4819 | valid |
| **Total:** | | | **12.432** | **85.471** | **12.41** | **2.48** | |

### Peptide QC Report 3811-13 58-79

| | |
|---|---|
| Analysis Name | D:\Data\3811-1358-79_267637_P1-C-3_01_141267.D |
| Sample Name | 3811-1358-79 D-Trp8-γ-MSH |
| Method | APRIL20171.2mLperMIN_NEPO A_141267.m |
| Instrument | amaZon SL |

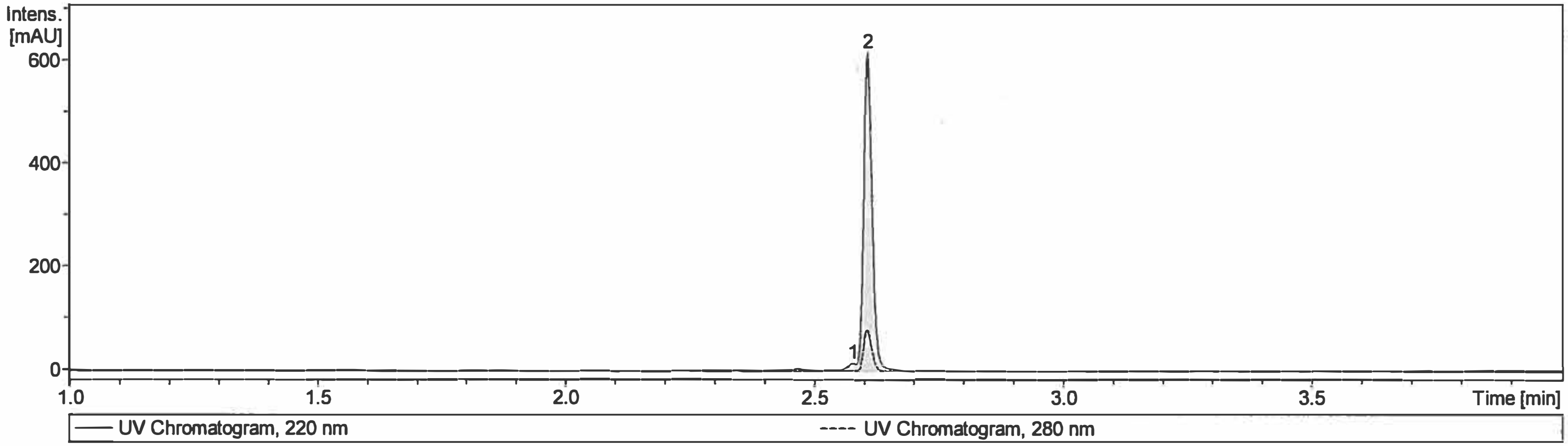


| Target Mass | Meas. Mass | Expec. Mass | Delt. Mr [Da] | Intensity | Area | Area Fraction [%] |
|---|---|---|---|---|---|---|
| Cmpd 2; 2.61 min; Pep Mr: 1570.70 | 1570.70 | 1570.80 | -0.10 | 612 | 729 | 97.2 |

| # | RT [min] | Area | Area Frac. % |
|---|---|---|---|
| 1 | 2.58 | 20.615 | 2.75 |
| 2 | 2.61 | 728.561 | 97.25 |

### Peptide QC Report 3811-22 11-44

| | |
|---|---|
| Analysis Name | D:\Data\3811-22 CYC 11-44_268411_P2-E-7_01_37512.d |
| Sample Name | 3811-22 11-44 Setmelanotide |
| Method | APRIL20171.2mLperMIN_NEPO A_37512.m |
| Instrument | UPLC1 |

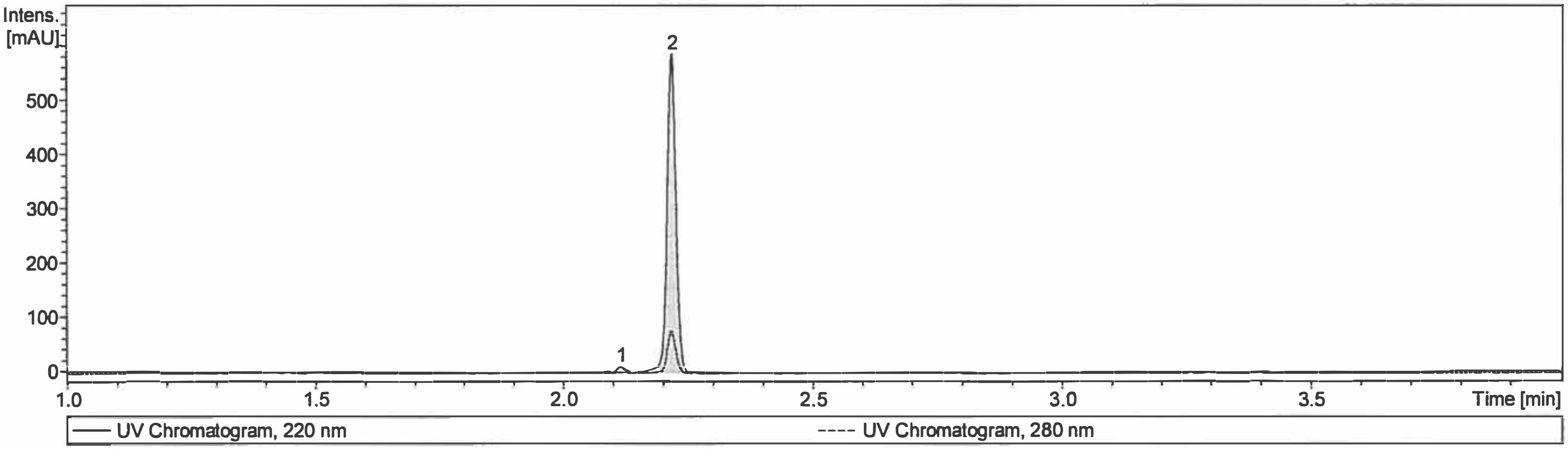


| Target Mass | Meas. Mass | Expec. Mass | Delt. Mr [Da] | Intensity | Area | Area Fraction [%] |
|---|---|---|---|---|---|---|
| Cmpd 2; 2.22 min; Pep Mr: 1117.68 | 1117.68 | 1117.30 | 0.38 | 584 | 706 | 98.5 |

| # | RT [min] | Area | Area Frac. % |
|---|---|---|---|
| 1 | 2.12 | 10.941 | 1.53 |
| 2 | 2.22 | 705.847 | 98.47 |

### Peptide QC Report BU04139 101-126

| | |
|---|---|
| Analysis Name | D:\Data\BU04139 101-126_279576_P1-C-8_01_42247.D |
| Sample Name | BU04139 101-126 SHU9119 |
| Method | APRIL20171.2mLperMIN_NEPOAHIGH_42247.m |
| Instrument | amaZon SL |

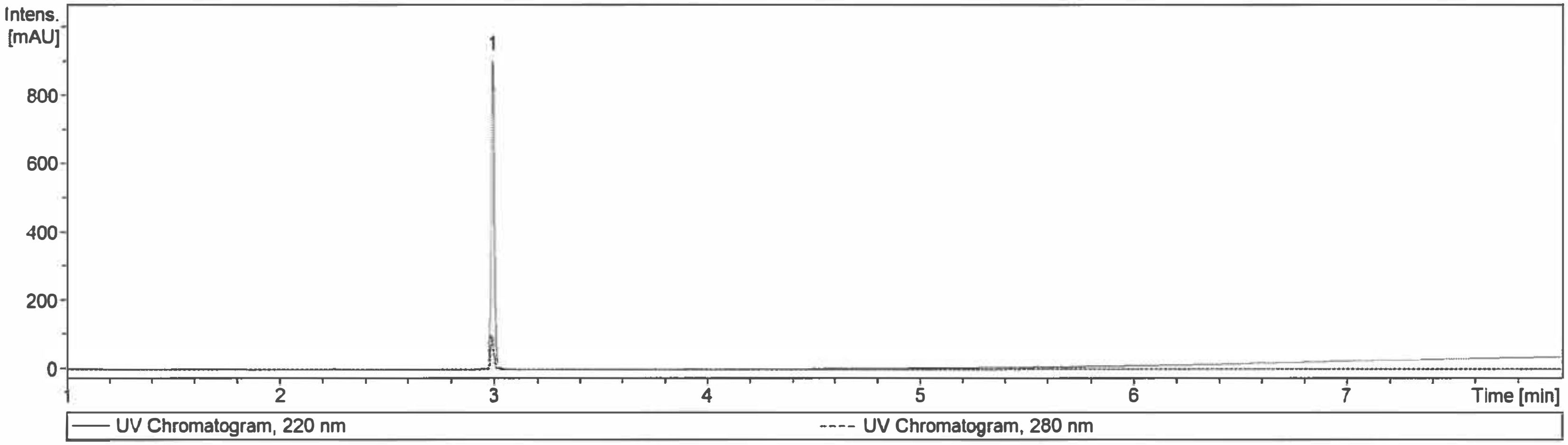


| Target Mass | Meas. Mass | Expec. Mass | Delt. Mr [Da] | Intensity | Area | Area Fraction [%] |
|---|---|---|---|---|---|---|
| Cmpd 1; 2.99 min; Pep Mr: 1074.55 | 1074.55 | 1074.10 | 0.45 | 915 | 1048 | 100.0 |

| # | RT [min] | Area | Area Frac. % |
|---|---|---|---|
| 1 | 2.99 | 1048.1 | 100.00 |

### Peptide QC Report 3811-11 47-64 Peptide CTX-1101

| | |
|---|---|
| Analysis Name | D:\Data\3811-1147-64_267677_P1-E-4_01_141286.d |
| Sample Name | 3811-11 47-64 |
| Method | APRIL20171.2mLperMIN_NEPO A_141286.m |
| Instrument | amaZon SL |

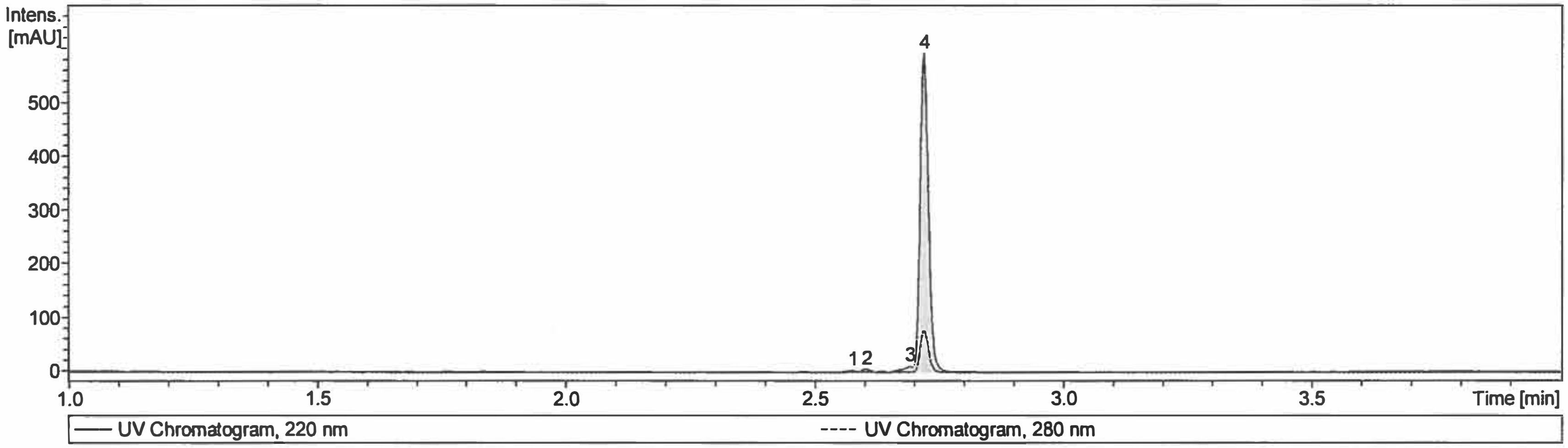


| Target Mass | Meas. Mass | Expec. Mass | Delt. Mr [Da] | Intensity | Area | Area Fraction [%] |
|---|---|---|---|---|---|---|
| Cmpd 4; 2.72 min; Pep Mr: 1552.84 | 1552.84 | 1552.70 | 0.14 | 589 | 688 | 97.6 |

| # | RT [min] | Area | Area Frac. % |
|---|---|---|---|
| 1 | 2.58 | 1.6910 | 0.24 |
| 2 | 2.60 | 3.9512 | 0.56 |
| 3 | 2.69 | 11.2777 | 1.60 |
| 4 | 2.72 | 688.4772 | 97.60 |

# Peptide QC Report    3891-29 32-65

| | |
|---|---|
| Analysis Name | D:\Data\3891-29 32-65_292972_P1-C-6_01_48068.D |
| Sample Name | 3891-29 32-65 CTX-2207 |
| Method | APRIL20171.2mLperMIN_NEPOAHIGH_48068.m |
| Instrument | amaZon SL |

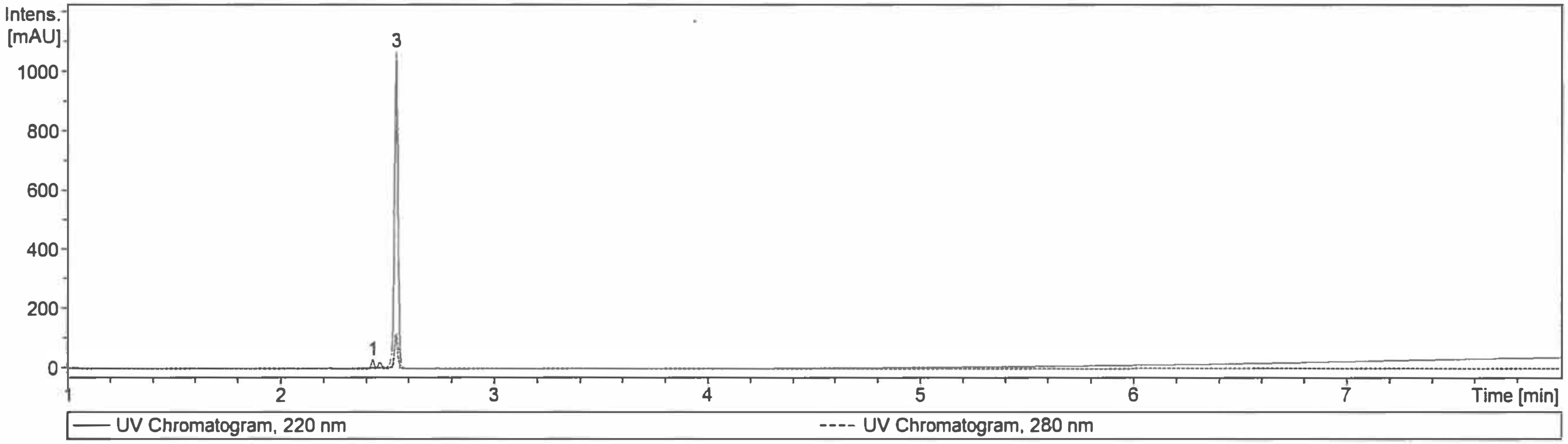


| Target Mass | Meas. Mass | Expec. Mass | Delt. Mr [Da] | Intensity | Area | Area Fraction [%] |
|---|---|---|---|---|---|---|
| Cmpd 3; 2.54 min; Pep Mr: 1167.59 | 1167.59 | 1167.30 | 0.29 | 1060 | 1253 | 96.7 |

| # | RT [min] | Area | Area Frac. % |
|---|---|---|---|
| 1 | 2.43 | 24.189 | 1.87 |
| 2 | 2.47 | 18.653 | 1.44 |
| 3 | 2.54 | 1252.558 | 96.69 |

### Peptide QC Report BU05301 31-55

| | |
|---|---|
| Analysis Name | D:\Data\BU05301 31-55_290809_P2-B-4_01_47083.D |
| Sample Name | BU05301 31-55 CTX-2100 |
| Method | APRIL20171.2mLperMIN_NEPO A_47083.m |
| Instrument | amaZon SL |

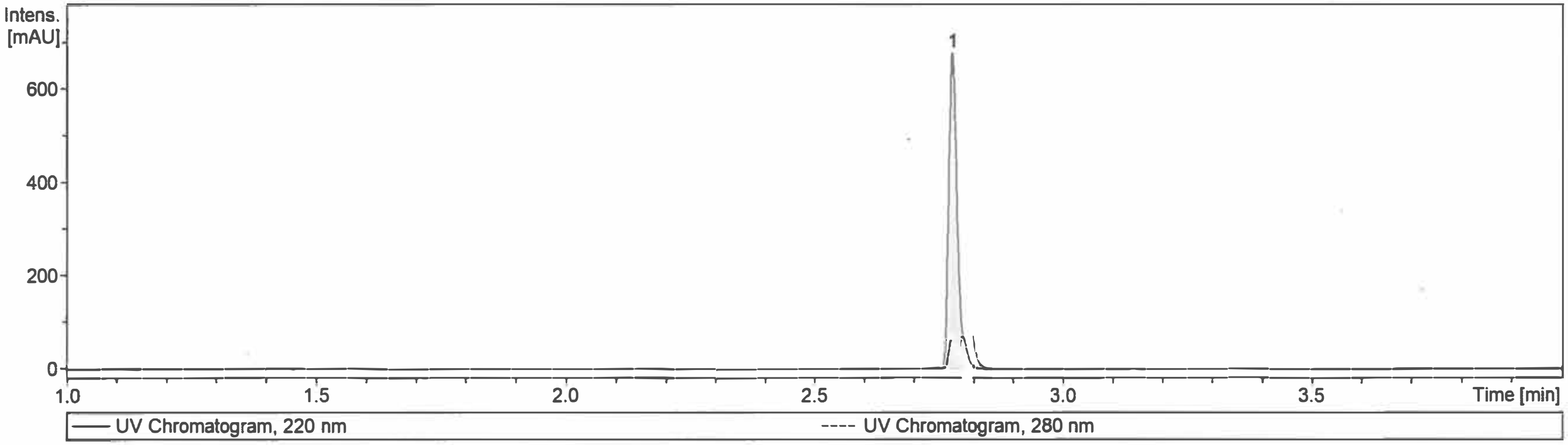


| Target Mass | Meas. Mass | Expec. Mass | Delt. Mr [Da] | Intensity | Area | Area Fraction [%] |
|---|---|---|---|---|---|---|
| Cmpd 1; 2.78 min; Pep Mr: 1602.81 | 1602.81 | 1602.60 | 0.21 | 675 | 796 | 100.0 |

| # | RT [min] | Area | Area Frac. % |
|---|---|---|---|
| 1 | 2.78 | 795.79 | 100.00 |

### Peptide QC Report BU03154 219-226

| | |
|---|---|
| Analysis Name | D:\Data\BU03154 219-226_273350_P1-C-9_01_144303.D |
| Sample Name | BU03154 219-226 CTX1306 |
| Method | APRIL20171.2mLperMIN_NEPOAHIGH_144303.m |
| Instrument | amaZon SL |

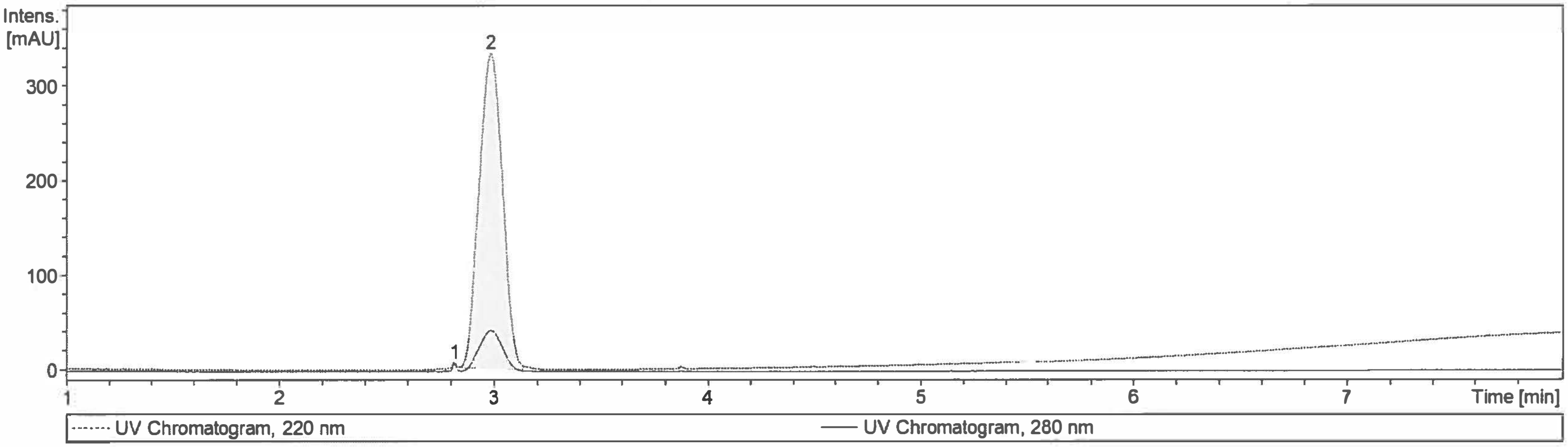


| Target Mass | Meas. Mass | Expec. Mass | Delt. Mr [Da] | Intensity | Area | Area Fraction [%] |
|---|---|---|---|---|---|---|
| Cmpd 2; 2.99 min; Pep Mr: 998.57 | 998.57 | 998.00 | 0.57 | 333 | 2503 | 99.7 |

| # | RT [min] | Area | Area Frac. % |
|---|---|---|---|
| 1 | 2.82 | 6.4206 | 0.26 |
| 2 | 2.99 | 2503.2029 | 99.74 |

### Peptide QC Report BU03200 214-223

| | |
|---|---|
| Analysis Name | D:\Data\BU032500214-223_273349_P1-C-8_01_144301.D |
| Sample Name | BU03200 214-223 cTX-2312 |
| Method | APRIL20171.2mLperMIN_NEPOAHIGH_144301.m |
| Instrument | amaZon SL |

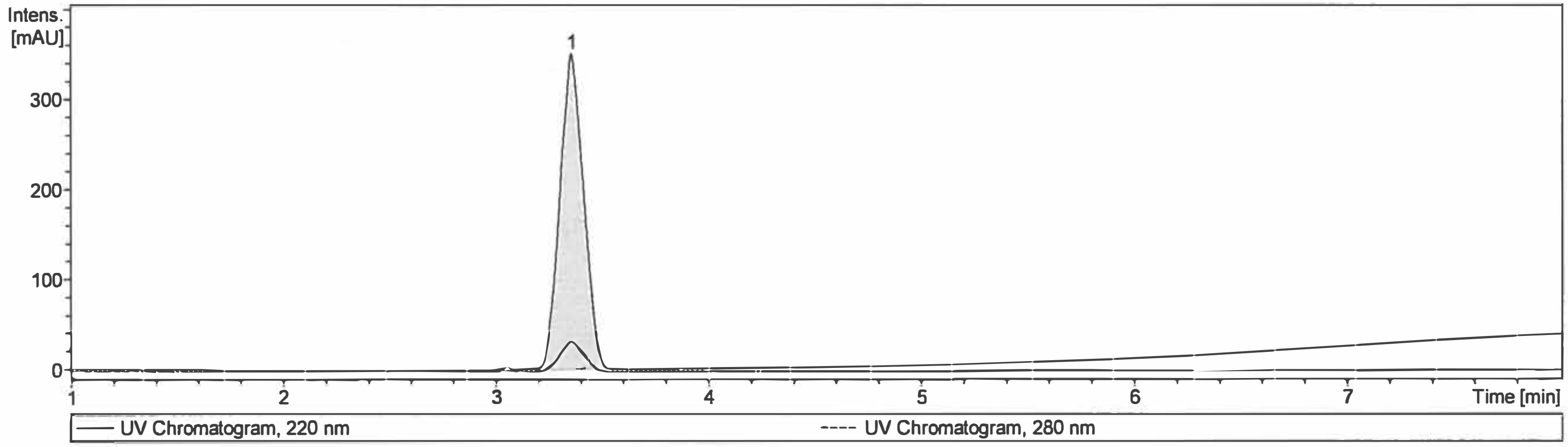


| Target Mass | Meas. Mass | Expec. Mass | Delt. Mr [Da] | Intensity | Area | Area Fraction [%] |
|---|---|---|---|---|---|---|
| Cmpd 1; 3.36 min; Pep Mr: 1048.59 | 1048.59 | 1048.10 | 0.49 | 350 | 2693 | 100.0 |

| # | RT [min] | Area | Area Frac. % |
|---|---|---|---|
| 1 | 3.36 | 2692.6 | 100.00 |

### Peptide QC Report 4162-34 21-50

| | |
|---|---|
| Analysis Name | D:\Data\NEP\4162-4 21-50_P1-E-2_1_7391.d |
| Sample Name | 4162-34 21-50 Analogue 11 |
| Method | NEPLC_NORM_7391.m |
| Instrument | UPLC1 |

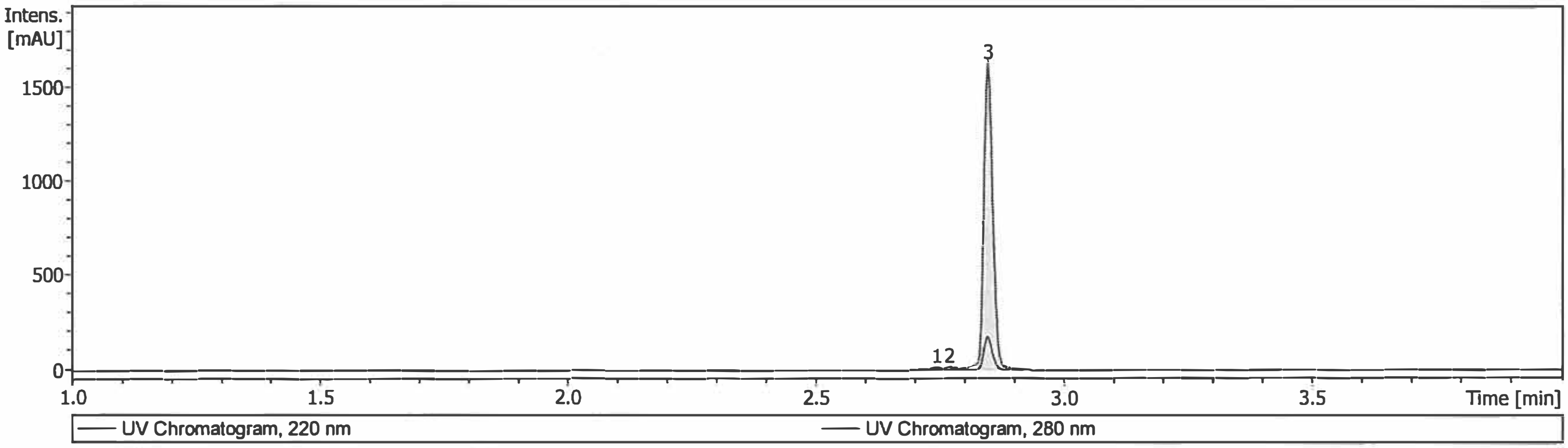


| Target Mass | Meas. Mass | Expec. Mass | Delt. Mr [Da] | Intensity | Area | Area Fraction [%] |
|---|---|---|---|---|---|---|
| Cmpd 3; 2.85 min; Pep Mr: 1601.77 | 1601.77 | 1601.60 | 0.17 | 1627 | 2011 | 99.2 |

| # | RT [min] | Area | Area Frac. % |
|---|---|---|---|
| 1 | 2.74 | 8.3700 | 0.41 |
| 2 | 2.77 | 7.4850 | 0.37 |
| 3 | 2.85 | 2010.7302 | 99.22 |

### Peptide QC Report 4162-32 31-73

| | |
|---|---|
| Analysis Name | D:\Data\NEP\4162-32 31-73_P1-E-3_1_7392.d |
| Sample Name | 4162-32 31-73 Analogue 13 |
| Method | NEPLC_HIGH_7392.m |
| Instrument | amaZon SL |

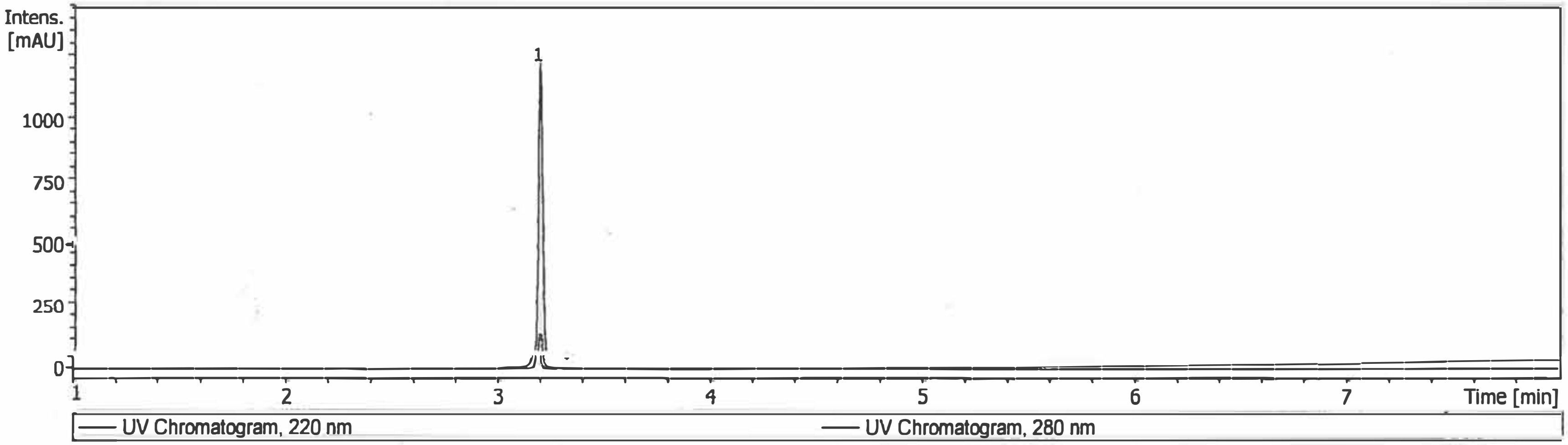


| Target Mass | Meas. Mass | Expec. Mass | Delt. Mr [Da] | Intensity | Area | Area Fraction [%] |
|---|---|---|---|---|---|---|
| Cmpd 1; 3.19 min; Pep Mr: 1561.74 | 1561.74 | 1561.60 | 0.14 | 1217 | 1773 | 100.0 |

| # | RT [min] | Area | Area Frac. % |
|---|---|---|---|
| 1 | 3.19 | 1772.6 | 100.00 |
